# Multidimensional Single-Nuclei RNA-Seq Reconstruction of Adipose Tissue Reveals Adipocyte Plasticity Underlying Thermogenic Response

**DOI:** 10.1101/2021.02.16.431320

**Authors:** Carlos Alberto Oliveira de Biagi, Sarah Santiloni Cury, Cleidson de Pádua Alves, Nabil Rabhi, Wilson Araujo Silva, Stephen R. Farmer, Robson Francisco Carvalho, Miguel Luiz Batista

## Abstract

Adipose tissue has been classified based on its morphology and function as white, brown, or beige / brite. It plays an essential role as a regulator of systemic metabolism through paracrine and endocrine signals. Recently, multiple adipocyte subtypes have been revealed using RNA sequencing technology, going beyond simply defined morphology but by their cellular origin, adaptation to metabolic stress, and plasticity. Here, we performed an in-depth analysis of publicly available single-nuclei RNAseq from adipose tissue and utilized a workflow template to characterize adipocyte plasticity, heterogeneity, and secretome profiles. The reanalyzed dataset led to the identification of different subtypes of adipocytes including three subpopulations of thermogenic adipocytes and provided a characterization of distinct transcriptional profiles along the adipocyte trajectory under thermogenic challenges. This study provides a useful resource for further investigations regarding mechanisms related to adipocyte plasticity and trans-differentiation.

**Highlights:** Multidimensional transcriptome analysis at single-nucleus resolution recovers nuclei of cell types in adipose tissue

Adaptative thermogenic response results in 3 distinct mature adipose cell types

Single-nuclei transcriptomic-based secretome analysis reveals adipose cell-type-specific genes

The *in vivo* trajectory of adipocyte plasticity for thermogenic response reveals sets of trans-differentiation genes

**Graphic Abstract:** 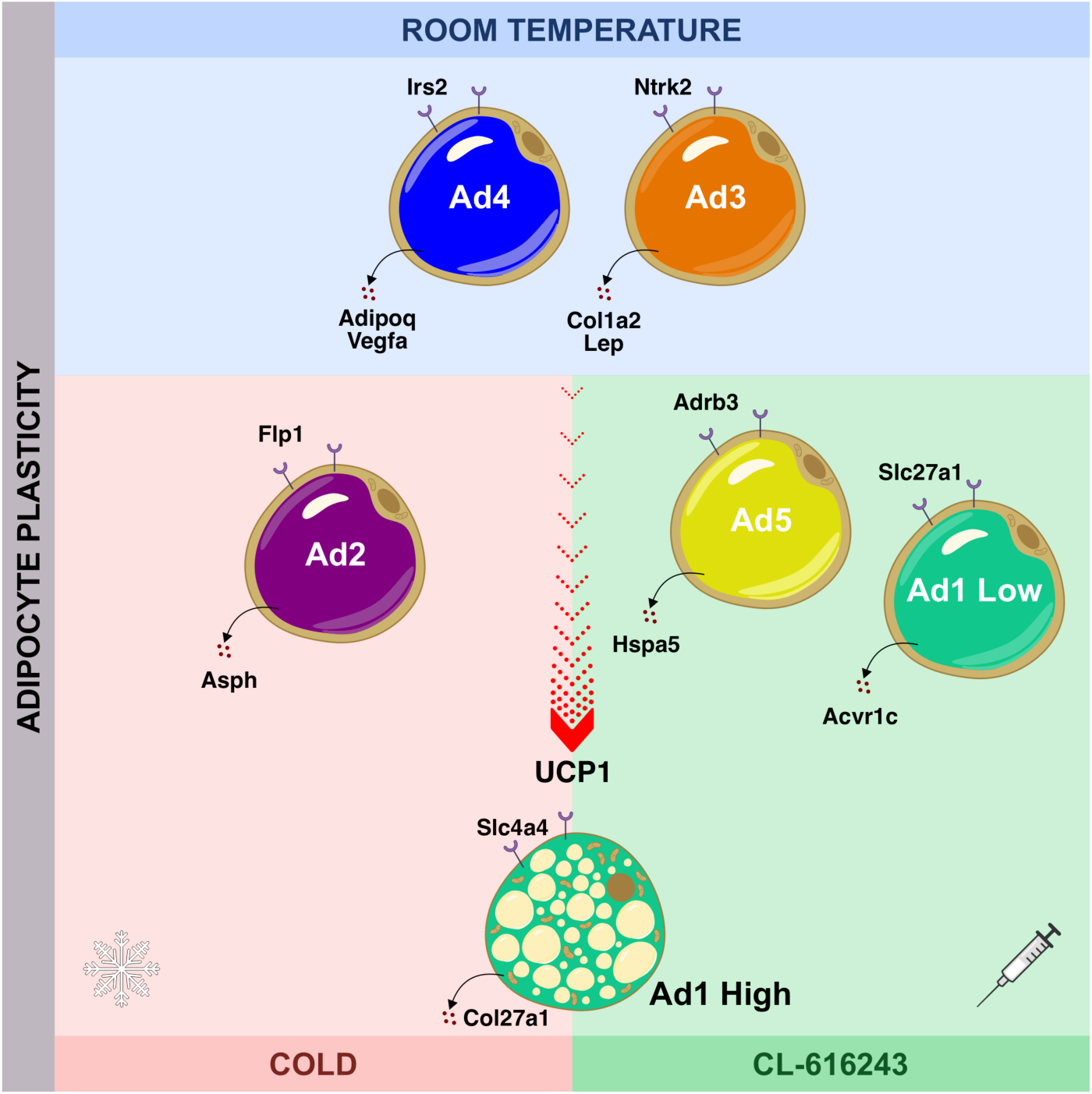

## Introduction

The perceived functional complexity of adipose tissue (AT) has changed significantly over the last 30 years since the leptin discovery (Friedman et al., 1991). The AT is a significant endocrine tissue organized into different depots, which are classified as brown (BAT) or white adipose tissue (WAT) (Scherer, 2006). Mature adipocytes constitute 90% of the AT volume but comprise only 17–33% of total cells. In contrast, the remaining vast majority of cells include a heterogeneous cell population of the stromal vascular fraction (SVF) (Lee et al., 2013a; Rosenwald and Wolfrum, 2014). Mature adipocytes are classified into three distinct types: white, brown, and beige/brite (Wang et al., 2016). White adipocytes are responsible for storing triacylglycerides (TGs). The brown adipocytes use lipids to produce heat in part through a UCP1 associated uncoupling of electron transport from ATP production (Pollard and Carling, 2020). Beige adipocytes (“brown-like”) can also support UCP1-independent thermogenesis (Pollard and Carling, 2020). These beige cells arise within white fat pads in response to activators such as cold exposure (Jiang et al., 2017), b3-adrenergic receptor (Adrb3) agonists (Lee et al., 2013b), PPARg ligands (Wang et al., 2016), cancer cachexia (Kir et al., 2014), and exercise training (Chouchani et al., 2019).

Beige adipocytes have been studied over the last three decades (Pollard and Carling, 2020). However, the interest in their physiological function and therapeutic potential to combat obesity has only recently been revisited after discovering the thermogenic response of white adipocytes in adult humans (de Jong et al., 2019; Wang et al., 2016). Formation of beige cells in WAT by cold or other stimuli occurs through *de novo* differentiation of progenitor cells from the perivascular compartment (Jiang et al., 2017) or interconversion of pre-existing white adipocytes (Jiang et al., 2017; Rosenwald et al., 2013).

Single-cell RNA sequencing (scRNA-Seq) has allowed for the identification of cell-to-cell heterogeneity and plasticity for many different tissues (Zheng et al., 2017; Ziegenhain et al., 2017). Analysis of adipose tissues at single-cell resolution is challenging. It has several limitations, especially considering technical limitations of reproducibly isolating the complete adipocyte compartment of the tissue due to the large size and high buoyancy of the adipocytes (Deutsch et al., 2020). Most WAT scRNA-Seq studies to date derive the transcriptomes of the cell types within SVF without providing critical information about the status of the adipocytes in animal models (Burl et al., 2018; Cho et al., 2019; Gu et al., 2019; Henriques et al., 2020; Hepler et al., 2018; Jaitin et al., 2019; Merrick et al., 2019; Schwalie et al., 2018; Spallanzani et al., 2019; The Tabula Muris Consortium et al., 2018; Weinstock et al., 2019; Zhang et al., 2019) and humans (Acosta et al., 2017; Jaitin et al., 2019; Merrick et al., 2019; Raajendiran et al., 2019; Vijay et al., 2020). Few studies investigated brown adipocytes (Song et al., 2019) or isolated adipocytes from mouse inguinal WAT undergoing browning (Rajbhandari et al., 2019). Most recently, Sárvári et al., investigated the complete repertoire of adipose tissue cell types at a single-cell resolution (Sárvári et al., 2021). However, multidimensional studies investigating the mechanisms involved in mature adipocyte plasticity under thermogenic stimuli are still lacking.

Intact cell nuclei have been used to perform single-nuclei RNA-seq (snRNA-seq), overcoming the limitations of isolating the complete adipocyte compartment (Nguyen et al., 2018). The snRNA-seq data of digested adipocytes from inguinal WAT reveals a complex subpopulation of mature adipocytes with distinct genetic signatures (Rajbhandari et al., 2019). Recently, snRNA-seq analysis of the WAT identified a rare subpopulation of adipocytes in mice that increase in abundance at higher temperatures. This subpopulation regulates the activity of neighboring adipocytes through acetate-mediated modulation of their thermogenic capacity (Sun et al., 2020).

Here, we reconstructed the mature adipocyte heterogeneity of the thermogenic response at the single-nuclei resolution. Our analyses generated a comprehensive and expansive cellular atlas presenting three thermogenic adipocyte subpopulations, followed by additional information on the metabolic pathways, the plasticity of individual subpopulations, and transcription factors possibly involved in beige remodeling of WAT. Also, we characterized specific cell surface markers and the secretome for each adipocyte subpopulation. The detailed snRNA-Seq analysis presented herein of the transcriptional changes in WAT adipocytes under thermogenic challenge provides insight into the molecular mechanisms driving adipocyte plasticity.

## Results

### Multidimensional snRNA-Seq reconstruction reveals distinct adipocyte subpopulations derived from mouse iWAT

We sought to characterize the transcriptional profiles of adipocytes by reanalyzing single-nuclei RNA sequencing data of isolated primary adipocytes responding to different thermogenic stimuli: 4°C challenge for four days (Cold) and CL-treatment, 1mg/kg/day for four days (CL). We selected the experimental challenges to increase the chance of detecting nuclei of mature primary adipocyte populations. For cells obtained from iWAT samples (from now, referred to as fat-cake), the t-SNE plots revealed 17 distinct nuclei clusters at different experimental conditions (**Figure S1A**). Those distinct clusters were subjected to a workflow template, depicted in **Figure 1A**. The first step of our pipeline was to subject raw data to over-clusterization (SCCAF), followed by the identification and classification of the cellular heterogeneity (MetaCell) (**Figure S1B**). We annotated the clusters of nuclei using marker genes (described in detail in the **STAR Methods**), which resulted in the identification of four groups of cell clusters: progenitor cells (PG), immune cells (IM), endothelial cells (EN), and adipocytes (AD1) (Fig. 1B). The proportion of cell types (average) per individual was 22% for progenitors, 10% for immune cells, 55% for endothelial cells, and 13% for adipocytes (**Figure S1B**). An accuracy threshold of 80% was used for cluster optimization (**Figure S1C**), and round 3 was chosen as the best round based on the accuracy and cross-validation test (**Figure S1D**). The distribution was similar after SCCAF over-clusterization: 22% for adipocyte progenitors and stem cells (PG1–PG5), 4% for immune cells (IM1–IM3) and 62% for endothelial cells (EN1–EN2), and 12% for adipocytes (AD1–AD4) (**Figure 1B**). Thus, unsupervised clustering of the single-nuclei transcriptional profiles identified four adipocyte subsets in the iWAT fat-cake. The following canonical cell type markers were upregulated in these clusters: *Pdgfra, Itgb1,* and *Cd34* (for adipocyte progenitors and stem cells), *Ptprc* (for immune cells), *Pecam1* (for endothelial cells), and Adrb3 (for adipocytes) (**Figure 1C**).

**Fig. 1.**
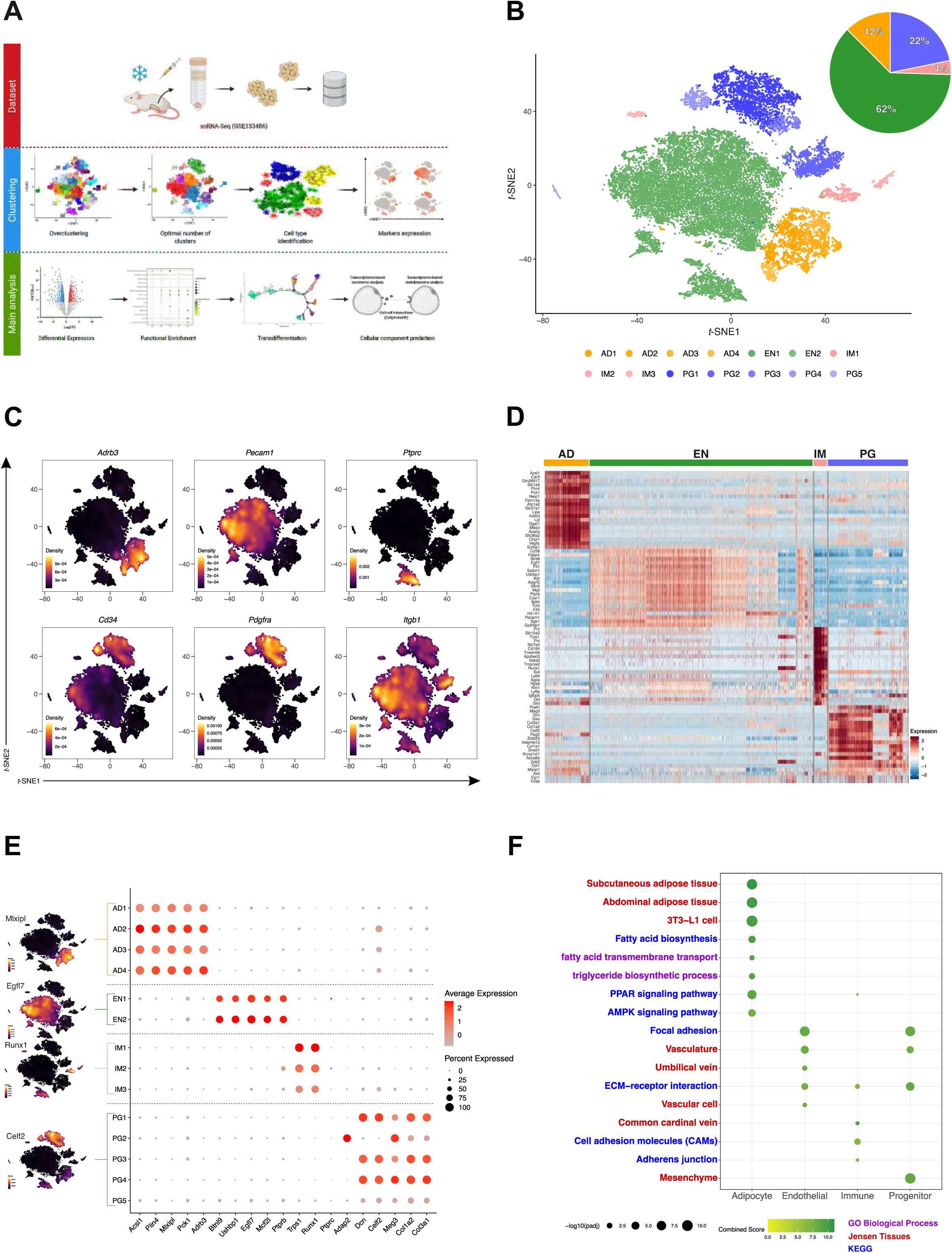
Adipose Single-Nuclei Workflow Template identifies different cell populations in adipose tissue. **(A)** Workflow overview showing the main 3 steps to process the template. The first step corresponds to obtaining the dataset (in this paper public data was used with accession number GSE133486). The next step is to cluster the data that includes the over clustering, finding the optimal number of clusters, cell type identification, and marker expression. The last step is related to the principal analysis that includes differential expression, functional enrichment, trans-differentiation, and cell component prediction. (Created with BioRender.com) **(B)** t-SNE plot of 28,820 single nuclei cells distributed by 14 clusters in 4 cell types: adipocyte (AD), endothelial (EN), immune (IM), and progenitors (PG). Pie charts show the corresponding percentage. **(C)** Nebulosa representing the unsupervised clustering of six canonical cell type markers for each cluster. *Pdgfra, Itgb1,* and *Cd34* (for adipocyte progenitors and stem cells), *Ptprc* (for immune cells), *Pecam1* (for endothelial cells), and Adrb3 (for mature primary adipocytes). **(D)** Gene-expression heatmap of the top 20 DEGs in each defined cell type compared to all others. Genes are represented in rows and cell clusters in columns. **(E)** Gene-expression dot plot of select top 5 DEGs for each defined cell type. Rows depict clusters, while columns depict genes. The intensity of any given point indicates average expression, while its size represents the proportion of cells expressing a particular gene. **(F)** Selected top categories from ORA analysis of DEGs from the four cell types identified. The intensity of the color in the dotplot indicates the enrichment significance by the combined score. Circle sizes correspond to the -log10 adjusted P-value (padj). Gene set names are colored according to the GO biological process (purple), Jensen tissues (red), and Kyoto Encyclopedia of Genes and Genomes (KEGG, blue).

After MetaCell analysis, a list of genes used to define each of the different clusters is presented in **Figure S1E** and **Table S1**. The expression profile of the top 20 cell-type-specific DEGs is shown in **Figure 1D** and **Table S2**. Unsupervised analysis of DEGs identified four significant adipocyte populations (*i.e*., expressing *Acsl1, Plin4, Mlxipl, Pck1,* and *Adrb3*), a population of endothelial cells (*Btnl9, Ushdp1, Egfl7, Ncf2l,* and *Ptprb*), adipocyte progenitors and stem cells (*Dcn, Celf2, Meg3, Col1a2,* and *Col3a1*), and immune cells (*Trps1, Runx1, Ptprc,* and *Adap2*) (**Figure 1E**). Adipocyte clusters enriched genes associated with subcutaneous adipose tissue and PPAR signaling. Endothelial cells enriched genes are related to focal adhesion and vasculature, immune cells are significantly enriched with cell adhesion molecules genes, and adipocyte progenitors and stem cells are enriched for mesenchyme cells and myofibroblasts genes (**Figure 1F**, **Table S3, Figure S1F**, and **Table S4**). These transcriptional differences may underlie distinct functional characteristics of the different cell types identified in the single-nuclei RNA-Seq reanalysis pipeline.

### Reclustering of adipocyte clusters reveals two distinct mature adipocyte populations at room temperature

To gain insight into the molecular differences between adipocyte subpopulations, we first verified the accuracy threshold of 80% that was used to cluster optimization (**Figure S2A**), and round 3 was chosen as the best round based on the accuracy and cross-validation test (**Figure S2B**). Next, we applied unsupervised over-clustering (SCCAF) to partition all 3,568 adipocytes nuclei that were identified using *t*-SNE. Interestingly, it identified five distinct adipocyte subpopulations (Ad1-Ad5, **Figure 2A**), each having a particular DEG pattern, with a slight exception for the Ad3 and Ad4 subpopulations (**Figure 2B** and **Table S5**). Canonical adipocyte markers *Dgat1, Plin1, Lipe, Cidec,* were expressed in all adipocyte subpopulations (**Figure 2C** and **Figure S2C**), albeit at varying levels. We found 571 DEGs in Ad1 (362 up and 209 down-regulated); 281 DEGs in Ad2 (118 up- and 163 down-regulated); 294 DEGs in Ad3 (70 up- and 224 down-regulated); 412 DEGs in Ad4 (353 up- and 59 down-regulated); and 160 DEGs in Ad5 (156 up- and 4 down-regulated) (**Figure S2D** and **Table S6**). We applied functional enrichment analysis of the five adipocyte subpopulations (Ad) (**Figure 2D**, **Table S7, Figure S2E,** and **Table S8**). The DEGs capture significant aspects of heterogeneity in distinct adipocyte subpopulations. Such differences were reflected in mitochondria gene expression and fatty acid degradation for Ad1, triglycerides biosynthetic process for Ad2, and ATP biosynthetic process for Ad1 and 2, TCA cycle and acetyl-CoA metabolic process, and regulation of cell differentiation for Ad3, regulation of sequestering triglycerides, and long-chain fatty acid transport, and adipocytokine signaling pathway for Ad4 and cholesterol metabolism for Ad5. Interestingly, fatty acid biosynthetic processes and long-chain fatty acid transport, and white adipose tissue (mouse-genes-atlas) were predominantly enriched in the Ad3 and Ad4 subpopulations. Interestingly, using the database Jensen tissues, we observed that the Ad3 subpopulation was the only one significantly enriched for the “Preadipocyte cell line” term (**Figure S2E**), suggesting a “preadipocyte-like” expression profile specifically found in the Ad3 (*Cfd, Fabp4, Gpd1,* and *Lpl*). Ad3 and Ad4 adipocyte subpopulations appeared to represent classical adipocytes, and they expressed genes associated with WAT, *Cidec, Pnpla,* and *Adipoq* (**Figure S2E**).

**Fig. 2.**
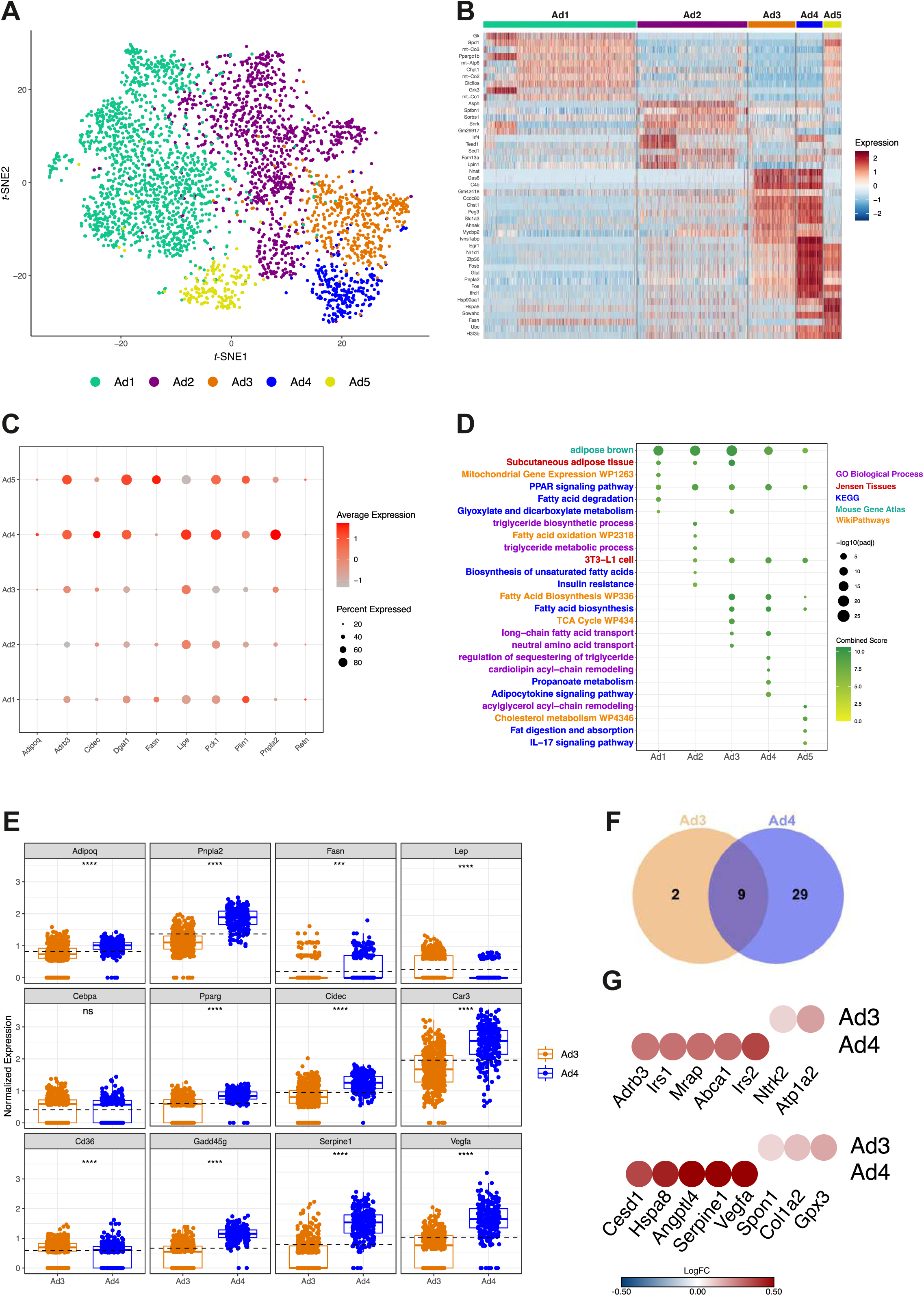
Multidimensional workflow template identifies two distinct mature adipocyte subclusters at RT. **(A)** Reclustering of adipocytes nuclei using SCCAF identified 5 putative adipocyte subclusters from “fat cake” iWAT nuclei data (Ad1 = 1,539 nuclei; Ad2 = 1,106 nuclei; Ad3 = 477 nuclei; Ad4 = 265 nuclei and Ad5 = 181 nuclei). **(B)** Gene-expression heatmap of the top 10 most DEGs in each adipocyte subcluster compared to all others. Genes are represented in rows and cell clusters in columns. **(C)** Gene-expression dot plot of the canonical adipocyte markers for each adipocyte subcluster. Rows depict clusters, while columns depict genes. The intensity of any given point indicates average expression, while its size represents the proportion of cells expressing a particular gene. **(D)** Selected top categories from ORA analysis of DEGs from the five adipocyte clusters. The intensity of the color in the dotplot indicates the enrichment significance by the combined score. Circle sizes correspond to the -log10 adjusted P-value (padj). Gene set names are colored according to the GO biological process (purple), Jensen tissues (red), Kyoto Encyclopedia of Genes and Genomes (KEGG, blue), Mouse Gene atlas (green), and WikiPathways (orange) . **(E)** Scatter-box plot representing normalized gene expression of 12 marker genes in the two adipocyte clusters from RT (Ad3 and Ad4 subclusters). At the top of each box is the significance of the comparison between Ad3 and Ad4 subclusters. The more stars, the more significant, while ns represents no significance. **(F)** Venn diagram demonstrating the intersection of upregulated genes encoding membrane proteins from Ad3 and Ad4 subclusters. **(G)** Heat-scatter plot representing two upregulated membranome components specifically identified in Ad3 adipocyte and the top five exclusive membranome of Ad4 subcluster (upper plot). Heat-scatter plot representing three upregulated secretome components specifically identified in Ad3 subcluster and the top five exclusive secretome of Ad4 subcluster (bottom plot). The color of the circles corresponds to the average logFC.

Ad3 and Ad4 subpopulations correspond to adipocytes present in fat cake of iWAT of non-treated mice (Control, RT) (**Figure 2A**). A comparison of adipocyte canonic markers revealed that Ad4 expresses higher levels of *Adipoq, Plpna2, Fasn, Pparg, Cidec, Car3, and Gadd45g* than Ad3 at room temperature (**Figure 2E**) suggesting that Ad4 more so than Ad3 consists of “classic’’ adipocytes. Interestingly, leptin is more highly expressed in Ad3 than in Ad4 subpopulation (**Figure 2E**).

To gain additional insight into Ad3 and Ad4 subsets, we performed an analysis of the secretome and membranome using gene expression profiles (DEGs). 29 genes predicted to encode membrane proteins were upregulated exclusively in Ad4, while only 2 genes were upregulated in Ad3 (*Ntrk2* and *Atp1a2*), and 9 other membrane genes were differentially expressed in both subpopulations (**Figure 2F**). **Figure 2G** shows the top 5 highly expressed genes, *Irs2, Abca1, Mrap, Irs1,* and *Adrb3*, which could potentially be used as Ad4 subpopulation markers. Regarding the secreted proteins, we found that Ad3 overexpresses 17.4% of genes that encode secretory proteins, while Ad4 overexpresses only 8.2% (**Figure S2F**). The top 5 exclusively expressed in Ad3 (*Gpx3, Col1a2*, and *Spon1*) and Ad4 (*Vegfa, Serpine1, Angptl4, Hspa8,* and *Cesd1*) subpopulation are detailed in **Figure 2G**. The complete list of predicted secretome and membranome upregulated genes from the Ad3 and Ad4 subpopulations is presented in **Figures S2G** and **Table S9**.

Characterization of the secretory proteins and components of the cell membrane permitted a prediction of cell-cell interactions via ligands and possible receptors (*i.e*., interactome) (**Figure S2H**). Ad3 and Ad4 showed increased interaction through collagens (produced by Ad3) and integrin (Ad4) and decreased interaction through NOTCH1 (Ad3) with JAG1 (Ad4) and COL5A1 (Ad3) and integrin complex (Ad4). Ad4 interacts with Ad3 by producing the ADIPOQ ligand interacting with the CLEC2D receptor in Ad3.

### Identification of a unique adipocyte thermogenic subpopulation corresponding to both Cold and CL-treatment

To further identify the adipocyte subpopulation with thermogenic transcriptome signature, we performed unbiased aggregated clustering of the processed data for each of the experimental conditions, *i.e.,* Cold, CL and RT, as a *t*-SNE-plot (**Figure 3A**). The aggregated cluster represents 3,027 adipocyte nuclei. **Figure 3B** shows the gene expression of the selected (supervised) adipocyte and thermogenic markers. For these data, it is interesting to note that the profile of gene distribution of *Adipoq, Retn, Cidec, and Fasn,* (canonical adipocyte markers), has almost no overlap with thermogenic genes, such as *Ppara, Ucp1, Dio2, Prdm16, Elovl3*. **Figure S3A** and **Table S10** highlight the DEGs related to cold-challenge (Cold) and CL treatment. The top 5 DEGs for cold-challenge were *Acacb, Acss2, mt-Co2, Macf1, and Gm26917,* while for CL were *Acsl1, mt-Co3, mt-Co2, mt-Atp6, Fasn.* Once we determined that the 2 treatments (Cold and CL) have different gene expression profiles, we performed functional enrichment analysis (**Figure 3C**, **Table S11**, **Table S12**, **Figure S3B,** and **Table S13**), using enriched genes in each cluster based on different experimental conditions. This analysis revealed that different adipocyte subclusters express distinct genes corresponding to the experimental conditions. For example, cold-induced subclusters demonstrated critical organophosphate biosynthetic processes and fatty acid transport. At the same time, CL showed significant aspects of organophosphate metabolic processes, oxidative phosphorylation, and ATP metabolic processes. The down-regulated genes reveal negative regulation for biosynthetic processes and response to mechanical stimuli for both experimental conditions.

**Fig. 3.**
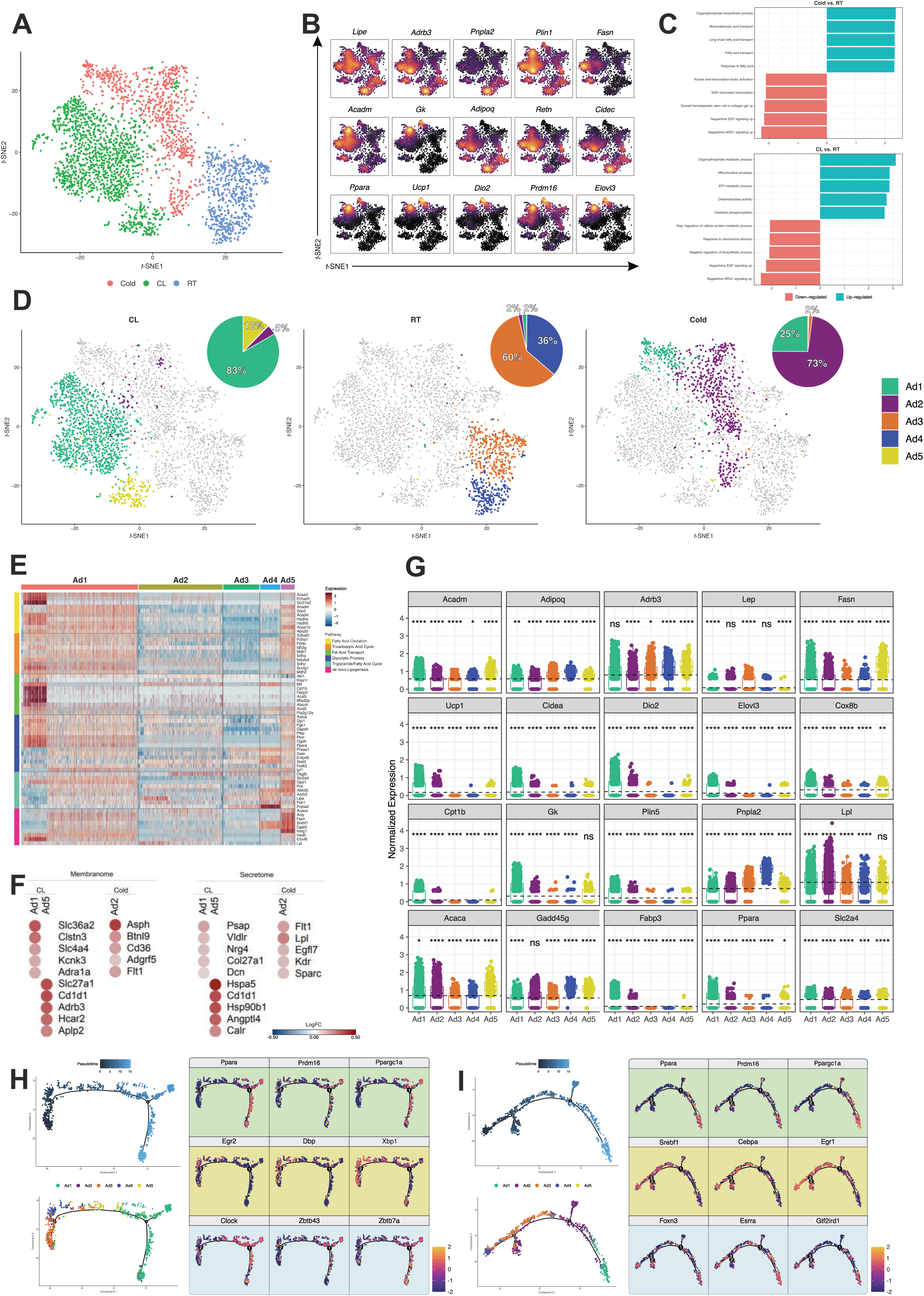
CL treatment and cold-challenge reveals a single thermogenic subpopulation. **(A)** t-SNE representation showing a subset of mature adipocyte nuclei subclusters classified according to each treatment (Cold = 892 cells: CL = 1,433 cells and RT = 702 cells). **(B)** Nebulosa expression plots of canonical mature adipocyte and thermogenic genes. **(C)** Gene Set Enrichment Analysis (GSEA) comparing Cold vs. RT and CL vs. RT. Rows depict enriched pathways, while in the y-axis, the Normalized Enrichment Score (NES). Positive NES represents an upregulated pathway, and negative NES represents down-regulated pathways. **(D)** CL, RT, and Cold shown subset of adipocyte nuclei subclusters classified according to each treatment integrated in t-SNE plot (CL = 1,443 cells; RT = 702 cells and Cold = 892 cells). Pie charts show the corresponding percentage. **(E)** Heat map showing the expression of main genes related to fatty acid oxidation, tricarboxylic acid cycle, fatty acid transport, glycolytic process, triglyceride/fatty acid cycle, and *de novo* lipogenesis in the five adipocytes subclusters. **(F)** Heat-scatter plot representing top five upregulated membranome components specifically identified in Ad1 subcluster and the top five exclusive membranome of Ad5 subcluster from CL subset comparison. The top five membrane genes from subcluster Ad2 (Cold) are demonstrated separately (left). Heat-scatter plot representing top five upregulated secretome components specifically identified in Ad1 subcluster and the top five exclusive secretome of Ad5 subcluster. The top five secretome genes from subcluster Ad2 (Cold) are demonstrated separately (right). The color of the circles corresponds to the average logFC. FC: fold change **(G)** Scatter-box plot representing normalized gene expression of 20 markers of canonical adipocyte and thermogenic genes in the five adipocyte subclusters. At the top of each box is the significance. The more stars, the more significant, while ns represents no significance. **(H)** Monocle-generated plots presenting pseudotime ordering and differentiation trajectory of CL and RT conditions. The trajectory suggests a transition between Ad3-Ad4-Ad5-Ad1. Green background represents the three main thermogenic TFs (classic). The yellow background represents the earlier expressed TFs in the trajectory, and the blue background represents the later expressed TFs in the trajectory. The characterized genes are DEGs throughout the trajectory. **(I)** Monocle-generated plots presenting pseudotime ordering and differentiation trajectory of Cold and RT conditions. The trajectory suggests a transition between Ad4-Ad3-Ad2-Ad1. Green background represents the three main thermogenic TFs (classic). The yellow background represents the earlier expressed TFs in the trajectory, and the blue background represents the later expressed TFs in the trajectory. The characterized genes are DEGs throughout the trajectory.

Since we have observed that RT, CL, and cold presents distinct frequencies in the adipocyte subpopulations, we analyzed the distribution of all adipocytes highlighted according to each different experimental condition (**Figure 3D**). Integrated analysis for adipocyte nuclei of CL, RT, and cold treatments revealed five subclusters, with Ad1 (83%) and Ad5 (12%) mainly from CL, Ad3 (60%) and Ad4 (36%) derived primarily from RT, and Ad2 (73%) and Ad1 (25%) mainly from cold. Interestingly, Ad1 contained adipocyte nuclei from both CL and Cold conditions and was more prevalent in Cl than other subpopulations. Whereas CL and Cold both exhibited unique expression patterns that reflected their functional commitments, CL showed significant functional enrichments mainly related to fatty acid degradation and transportation (**Figure S3B**). Unsurprisingly, the Ad1 subpopulation was most heavily involved in fatty acid oxidation, TAC, and fatty acid transport (**Figure 3E** and **Figure S3C**). The latter seems to be very specific to Ad1. The expression profiles of genes corresponding to glycolytic process and triglycerides / fatty acid cycle, however, were not so distinguished between Ad1 *versus* Ad5 subpopulation (**Figure 3E**), despite the fact the Ad5 showed relatively higher expression of most of the metabolism-related genes, in particular, triglycerides / fatty acid processes and *de novo* lipogenesis; this is indicative of a high energy demand for Ad5 subpopulation.

Identification of three distinct thermogenic adipocyte subpopulations led us to predict corresponding secretome and membranome (**Figure 3F** and **Figures S3D-S3G,** and **Table S14**). In Ad1 and Ad5, 7.2 % and 9%, respectively, of upregulated genes represent membrane encoding proteins (**Figure S3E**). Two genes were exclusive to both Ad1 and Ad5 clusters (**Figure S3D**), while 23 plasma membrane genes were upregulated exclusively in Ad1 and 12 genes in Ad5; the top 5 expressed genes in each subpopulation are *Slc36a2, Clstn3, Slc4a4, Kcnk3, Adra1a* for Ad1, and *Slc27a1, Cd1d1, Adrb3, Hcar2, Aplp2 fo*r Ad5 (**Figure 3F**). The Ad1 subcluster was found to overexpress 3.3% of potential secreted proteins and Ad5 overexpressed 5.8%. The secretory genes exclusively expressed in Ad1 and Ad5 subpopulations were: Ad1: *Psap, Vldlr, Nrg4, Col27a1*, *Dcn;* Ad5*: Hpsa5, Cd1d1*, *Hsp90b1, Angptl4, Calr* (**Figure 3F**). The complete list of the predicted secreted and membrane proteins in Ad1 and Ad5 subpopulations are presented in **Figure S3F and G**. Ad2 shows 12.7 % of upregulated genes are membrane encoding proteins (*Asph, Btnl9, Cd36, Adgrf5,* and *Flt1*) and 9.3% are secreted proteins (*Flt1, Lpl, Egfl7, Kdr,* and *Sparc*). VEGFA was the main interacting molecule between Ad1 and Ad5 subclusters (**Figure S3H**).

The gene expression profile (normalized) of the adipocyte subpopulations shows that the classic thermogenic markers, *Ucp1, Cidea, Dio2, Elovl3, Cpt1b,* and *Plin5*, are differentially expressed in all clusters, but particularly in Ad1 (**Figure 3G**). Other genes, such as *Acadm, Cox8b,* and *Gk* seem to be closely related to CL subclusters (Ad1 and Ad5). *Fasn* and *Acaca* showed a higher level of gene expression in Ad5 than all other subpopulations, which may indicate a specific profile in this adipocyte phenotype.

Since reanalysis of the snRNA-seq dataset simultaneously profiles adipocytes under both cold and CL-treatment, we hypothesized that their plasticity could be traced *in vivo* by mapping a developmental trajectory between distinct adipocyte subpopulations. As two significant subpopulations were resolved in the RT nuclei dataset (Ad3, Ad4) from iWAT (**Figure 3D**), we established this condition as the starting point for the analysis through the Monocle. Pseudotime study mapped a distinct trajectory of RT adipocytes responding to thermogenic challenge into different cellular states, ranging over modifications to late thermogenic trans-differentiation (**Figures 3H** and **3I**). This identified a branched trajectory connecting the adipocyte subpopulations with two branches representing the adipocytes’ specification into distinct subpopulations. We focused on the thermogenic branch 1 to separate the nuclei into late trans-differentiation (mostly Ad1), and in branch 2, related to early trans-differentiation (mostly Ad3) (**Figure S3I and S3J**).

In the CL subpopulations (Ad1, Ad5), adipocytes lost expression of transcription factors (TF) *Egr2, Dpb,* and *Xbp1*, as well as of genes *Nnat, Plppr3,* and *Chst1* (**Figure S3I**). A progressive gain of expression is shown for TFs, *Clock, Zbtb43,* and *Zbtb7a*, whereas “classic” thermogenic TFs *Ppara, Prdm16,* and *Ppargc1a* are expressed along the entire trajectory analysis. *Ucp1* was expressed in late trans-differentiation thermogenic adipocytes, followed by *Ppara* and *Dio2* (**Figure S3I** and **Table S15**). The cold-challenged adipocytes (Ad1 and Ad2) lost expression of *Egr1, Cebpa,* and *Srebf1* and progressively gained expression of TFs *Foxn3, Essra*, and *Gtf2ird1* (**Figure 3I**). Brown adipocyte markers were also expressed in late trans-differentiation thermogenic adipocytes (**Figure S3J** and **Table S16**).

### Ad1-*Ucp1*^High^ subpopulation shows a “classical” thermogenic profile

Once the Ad1 subpopulation was shown to express a clear thermogenic signature in response to both Cold- and CL-challenge, the next step was to evaluate Ad1 nuclei that express high levels of UCP1 and compare them with those expressing low levels of UCP1 **(details in STAR Method**). **Figure 4A** presents the *t*-SNE representation of adipocyte subpopulations Ad1, Ad2, and Ad5 highlighting nuclei with high expression of *Ucp1* (from hereafter, Ad1 will be named Ad1-*Ucp1*^High^ and *Ucp1*^Low^). The thermogenic genes *Ppara, Dio2, Prdm16, Elovl3, Cox8b* are more prevalent in the Ad1 subpopulation (**Figure S4A**). The top-5 DEGs in Ad1-*Ucp1*^High^ nuclei are *Macf1, Gk, Grk3, Pdk4*, and *Acacb* (**Figure 4B**, **Table S17, Figure S4B,** and **Table S18**). Interestingly, it should be noted that a few adipocyte nuclei belonging to Ad1-*Ucp1*^Low^ have a gene signature similar to Ad1-*Ucp1*^High^.

**Fig. 4.**
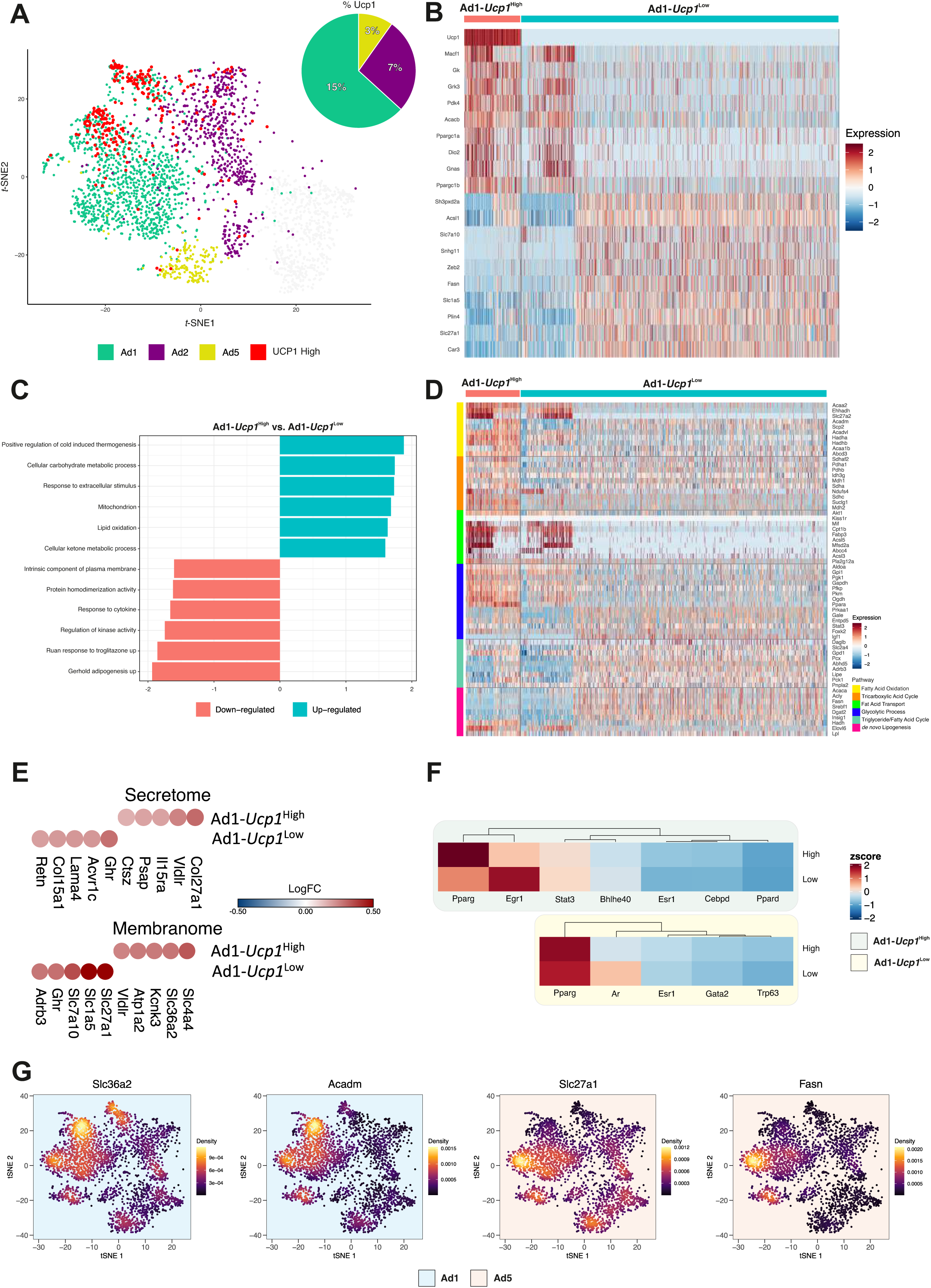
Ad1-*Ucp1*^High^ subpopulation shows a “classical” thermogenic profile whereas Ad1-*Ucp1*^Low^ subpopulation suggest a potential UCP1-independent pathways activation. **(A)** *t*-SNE representation showing Ad1, Ad2 and Ad5 mature adipocyte nuclei subclusters. The red dots represent the cells that have higher expression levels of *Ucp1*. Pie charts show the corresponding percentage of high *Ucp1* in each of Ad1, Ad2 and Ad5 subclusters. **(B)** Gene-expression heatmap of the top 10 most DEGs in Ad1-*Ucp1*^High^ vs. Ad1-*Ucp1*^Low^ comparison. Genes are represented in rows and cell clusters in columns. **(C)** Gene Set Enrichment Analysis (GSEA) comparing Ad1-*Ucp1*^High^ vs. Ad1-*Ucp1*^Low^. Rows depict enriched pathways, while in y-axis the Normalized Enrichment Score (NES). Positive NES represents an upregulated pathway and negative NES represents down-regulated pathways. **(D)** Heat map showing the expression of main genes related to fatty acid oxidation, tricarboxylic acid cycle, fatty acid transport, glycolytic process, triglyceride/fatty acid cycle, and *de novo* lipogenesis in the Ad1-*Ucp1*^High^ and Ad1-*Ucp1*^Low^. **(E)** Heat-scatter plot representing the top five upregulated secretome components identified in Ad1-*Ucp1*^High^ and Ad1-*Ucp1*^Low^ (upper plot). Heat-scatter plot representing the top five upregulated membranome components identified in Ad1-*Ucp1*^High^ and Ad1-*Ucp1*^Low^ (bottom plot). The color of the circles corresponds to the average logFC. **(F)** Heatmap showing the average expression for seven main TFs in Ad1-*Ucp1*^High^ cells (green background) and five main TFs in Ad1-*Ucp1*^Low^ cells (yellow background). These TFs are related to the DEGs from each comparison through the Transcription Factor Enrichment Analysis (TFEA) using the X2K database. **(G)** Nebulosa expression plots representing four positive/negative markers for *Ucp1* (*Slc36a2, Acadm, Slc27a1* and *Fasn*).

To gain mechanistic insight into gene lists, we applied enrichment analysis to evaluate the pathways enriched in a gene list of Ad1 (*Ucp1*^High^ and *Ucp1*^Low^). As expected, for Ad1-*Ucp1*^High^, pathways related to positive regulation of cold-induced thermogenesis, mitochondrion, lipid oxidation, and response to extracellular stimulus are activated (**Figure 4C**). For Ad1-*Ucp1*^Low^, pathways related to adipogenesis, response to troglitazone, and regulation of kinase activity showed more activation.

To gain additional information about the central metabolic pathways activated in the Ad1-*Ucp1*^High^ and Ad1-*Ucp1*^Low^, we analyzed the DEGs of the primary genes involved in distinct energy metabolism pathways (**Figure 4D** and **Figure S4C**). The group of genes related to TCA is preferentially activated in Ad1-*Ucp1*^High^ over Ad1-*Ucp1*^Low^. This profile was followed by the upregulation of genes involved in the following pathways: fatty acid oxidation, fatty acid transport, and glycolytic processes. Interestingly, Ad1-*Ucp1*^Low^ has a higher enrichment of triglycerides / fatty acid cycle genes than in Ad1-*Ucp1*^High^. This fact suggests that other metabolic pathways, particularly those involved in a futile cycle, could be affected (**Figure 4D** and **Table S19**). Aware of this fact, we also evaluated non-canonical or UCP1 independent pathways involved in the thermogenic program, such as glycolytic pathway, creatine metabolism genes, such as *Gamt*, *Gatm,* and *Ckmt1*, and SERCA2-pathway, such as *Arpc2, Adra1a, Atp2a2,* and *Tmlc4* (**Figure S4C**). Overall, there were no categorical differences in these programs when comparing Ad1-*Ucp1*^High^ and Ad1*-Ucp1*^Low^. However, concerning the glycolytic pathway, *Ppara*, *Pkm*, and *Ogdh*, particularly the former, are differentially expressed in Ad1-*Ucp1*^High^, while *Atp2a2* is equally represented in Ad1-*Ucp1*^High^ and Ad1-*Ucp1*^Low^ adipocytes. For Ad1-*Ucp1*^Low^, triglycerides / fatty acid processes and *de novo* lipogenesis are more enriched, with highlights for the genes; *Fasn*, *Srebf1* and *Insig1*.

Thirty-seven membrane genes were upregulated exclusively in Ad1-*Ucp1*^High^ and 14 genes upregulated in Ad1-*Ucp1*^Low^ (**Figure S4D** and **Table S20**). The top 5 highly expressed genes are *Slc4a4, Slc36a2, Kcnk3, Atp1a2, and Vldlr* for Ad1-*Ucp1*^High^, and *Slc27a1, Slc1a5, Slc7a10, Ghr, and Adrb3* for Ad1-*Ucp1*^Low^ (**Figure 4E**). The top 5 genes predicted to encode secreted proteins for Ad1-*Ucp1*^High^ were *Col27a1*, *Vldlr*, *Il15ra*, *Psap*, and *Ctsz,* while for Ad1-*Ucp1*^Low^ were *Ghr*, *Acvr1c*, *Lama4*, *Col15a1*, and *Retn* (**Figure 4E**).

Given that TFs are crucial for defining cell identity, we analyzed TF expression levels in the Ad1 subpopulation. For this analysis, we initially evaluated the TFs that are most likely involved in regulating DEGs for Ad1-*Ucp1*^High^ and Ad1-*Ucp1*^Low^ (**Figures S4E** and **S4F**). The former exhibited specific expression of the main canonical adipocyte TFs (such as *Parpd, Hnf4a, Ers1, Pparg, Sall4, Cebpd, Egr1, Nanog, Stat3, and Bhlhe40*). Protein-protein interactions (PPI) analysis demonstrated that *Arpd, Hnf4a, Ers1, Pparg, Sall4, Egr1, Nanog*, and *Bhlhe40* could interact with the targeted genes differentially expressed in *Ad2-Ucp1^High^*. For Ad1-*Ucp1*^Low^, *Parpg, Sall4, Ers1, Tp63, Ar,* and *Gata2*, the most enriched TFs and PPI analysis demonstrated that *Parpg, Ers1, Tp63,* and *Gata2* could interact with the targeted DEGs. The next step was to analyze the most highly expressed TF genes (**Figure 4F**), in addition to exploring the genes that are targets of these TFs (**Figure S4G**). As expected, the canonical adipocyte TFs, such as *Pparg*, are expressed at higher levels than the other TFs in both Ad1-*Ucp1*^High^ and Ad1-*Ucp1*^Low^, being more intense in the former. On the other hand, *Egr1* and *Ar* are more pronounced in Ad1-*Ucp1*^Low^ adipocytes. The target genes related to TFs, *Slc4a4, Slc25a42*, and *Pdk4,* were more pronounced in Ad1-*Ucp1*^High^ adipocytes while *Slc1a5* and *Fasn* were upregulated in Ad1-*Ucp1^L^*^ow^ compared to Ad1-*Ucp1*^High^. **Figure 4G** shows the intracellular and membrane markers for Ad1-*Ucp1*^High^ are *Slc36a2 and Acadm),* while *Slc27a1* and *Fasn* correspond to Ad1-*Ucp1*^Low^.

## Discussion

This report analyzed existing single-nuclei transcriptome data to gain a greater understanding of the cellular composition of WAT. Here, we generate a comprehensive cellular atlas and classification of the adipocytes into five distinct subpopulations. This allowed us to delineate *in vivo* trajectories and determine the plasticity of individual adipocyte subpopulations from the inguinal fat pad of mice in the setting of thermogenic challenges. Cold and CL both induce UCP1+ cell populations (Ad1-*Ucp1*^High^) in addition to two other adipocyte subpopulations that show gene expression signatures distinct from each other. We also identify a new adipocyte population (Ad5) specific to CL treatment, which demonstrates enrichment for lipid turnover and *de novo* lipogenesis pathways, suggesting this subpopulation provides a higher energy output in a UCP1-independent fashion. We further show that the different adipocyte subpopulations present specific secretome profiles, mainly composed of proteins secreted via classical and non-classical pathways (such as exosomes).

In WAT, beige remodeling can be triggered through two stimuli: Cold and CL (Jiang et al., 2017; Rosenwald et al., 2013). Recent studies suggest that these two stimuli might induce beiging through distinct pathways and our data support those observations (Jiang et al., 2017; Rabhi et al., 2020). The CL-induced subpopulation showed significant oxidative phosphorylation and ATP metabolic processes, whereas cold-induced thermogenic adipocytes are more specialized for fatty acid transport. As expected, a unique mature adipocyte subpopulation (Ad1) showing a thermogenic “classic” signature, such as *Ucp1, Cidea, Dio2, Elovl3, Cpt1b,* and *Plin5*, was detected in both treatments. The thermogenic population showed to be specialized in the following pathways: fat acid oxidation, TAC, and fatty acid transport.

Interestingly, CL treatment also resulted in an additional subpopulation, Ad5. Despite not having a classic thermogenic signature, this adipocyte subpopulation showed a profile for activating genes related to glycolytic, fatty acid turnover, and, in particular, *de novo* lipogenesis pathways, also present at some level in thermogenic Ad1. Ad5 also showed specialization in lipid turnover pathways, suggesting this subpopulation provides a higher energy output. *Acadm, Cox8b,* and *Gk* are associated with both Ad1 and Ad5 subpopulations, while *Fasn, Acly,* and *Insig1* is more highly associated with the Ad5 subpopulation. This set of results, predominantly generated *in silico*, meets the *ex-vivo* analysis presented by Lee et al., 2017. In this paper, Granneman and colleagues showed that CL upregulated FASN and MCAD in distinct adipocyte populations: high MCAD expression in multilocular adipocytes that co-expressed high UCP1 levels, while FASN expression occurred in paucilocular adipocytes with low UCP1 levels. These results corroborate the concept of metabolic heterogeneity as a distinct property of activated thermogenic adipocytes. However, the function of each population and the control mechanisms involved need further analysis.

Regarding the secretome of thermogenic subpopulations, Ad1 adipocytes differentially overexpressed *Nrg4, Col27a1*, *Psap* and *Dcn,* whereas the Ad5 subcluster over-expressed mainly *Hpsa5*, *Hsp90b1* and *Cd1d1*. Neuregulin 4 (NRG4) was previously shown as an AT-enriched secreted factor, markedly increased during brown adipocyte differentiation. *Nrg4* is downregulated in AT during rodent and human obesity. In contrast, gain-and-loss-of-function studies in mice showed it protects against diet-induced insulin resistance and hepatic steatosis (Wang et al., 2014). Interestingly, NRG4 promotes neurite outgrowth during cold-challenge (Rosell et al., 2014). CD1d is a lipid antigen-presenting molecule for iNKT cells (invariant Natural Killer T), highly expressed in adipocytes than any other cell types of adipose tissue (Hotamisligil et al., 1993). Adipocytes positively express CD1d molecules in lean adipose tissue, which play a crucial role in maintaining the adipose iNKT cell population. Upon HFD feeding, adipocytes present obesity-related lipid antigens via CDld molecules, which leads to iNKT cell activation and stimulates anti-inflammatory cytokine secretion from adipose iNKT cells (Huh et al., 2018). In this sense, the adipocyte secretome prediction has shown that both NRG4 (Ad1) and CD1d (Ad5) are specific thermogenic adipocyte products since sufficient mRNA levels molecules were not detected in other cell populations from WAT.

To fully understand the potential therapeutic relevance of beige remodeling, it is crucial to characterize the overall metabolic properties of beige adipocytes in addition to their thermogenic potential. In this regard, the Ad1 subpopulation was subdivided into two subpopulations, according to the presence or absence of *Ucp1* mRNA. Interestingly, although Ad1-*Ucp1*^High^ showed a “canonical” thermogenesis genetic signature, by expressing higher DEGs corresponding to energy metabolism, particularly those related to TCA, Ad1-*Ucp1*^Low^ adipocytes showed higher triglycerides/fat acid cycle and *de novo* lipogenesis pathways. Also, specialization of fatty acid metabolism (oxidation and transport) and glycolytic processes are more pronounced in Ad1-*Ucp1*^High^ despite a small portion of Ad1-*Ucp1*^Low^ adipocytes having a similar profile to that found in Ad1-*Ucp1*^High^. Ad1-*Ucp1*^High^ showed, in addition to the positive regulation of thermogenesis, specialization related to response to extracellular stimulus and cellular carbohydrate metabolic process. Our analyses also showed that the UCP1 independent pathways involved in the thermogenic program, such as glycolytic, and SERCA2-pathway are equally expressed in Ad1-*Ucp1*^High^ and Ad1*-Ucp1*^Low^ adipocytes. This fact suggests that other metabolic pathways, particularly those involved in a futile cycle (lipid turnover and SERCA2), could also be involved in the thermogenic metabolism in Ad1 adipocytes in a way that does not depend on the presence or absence of UCP1. Furthermore, it should be highlighted that Ad1*-Ucp1*^Low^ adipocytes make up about 75% of the adipocyte subpopulation under CL treatment, suggesting its relevance for the detailed understanding of beige remodeling.

In recent years, beige remodeling has been demonstrated through the existence of other thermogenic pathways in a UCP1-independent manner (Kajimura et al., 2015; Szabo and Zoratti, 2017). Even more recently, Kajimura and colleagues showed ‘glycolytic beige’ (g-beige), with significant enrichment of genes involved in glycolysis, glucose, and carbohydrate metabolism distinct from both the classical beige and brown adipose signatures (Chen et al., 2019). However, at present, further analysis is necessary to gain insights into whether these new cell types are expressed in wild-type mice or if their expression requires specific genetic backgrounds.

These findings motivated us to identify new markers for thermogenic adipocyte types using snRNA-seq data from digested AT samples. Although some consistent studies have presented the transcriptome profile of beige cells, either of bulk (Lee et al., 2013b) or cells isolated by a reporter system (UCP1+) (Roh et al., 2018), as far as we know, this is the first analysis of the snRNA-Seq data to predict the profile of proteins associated with their respective functions. The membranome prediction analysis showed *Slc4a4* and *Adra1a* and *Slc36a2* as specific for Ad1-*Ucp1*^High^ surface markers, whereas *Slc27a1* and *Slc1a5* were more typical for Ad1-*Ucp1*^Low^. Our analyses show a very particular profile of surface proteins (plasmatic membrane), making possible additional functional analyses of those adipocyte populations. However, experimentally using *ex vivo* and in *vivo* or approaches such as sorting cells (Roh et al., 2018) and fluorescent imaging techniques must be determined.

Brown/beige adipocyte differentiation and activation of the thermogenic program are controlled by sequential actions of transcription factors (TFs), including EBF2, PRDM16, C/EBPΒ, PGC-1α, and PPARγ (Kajimura et al., 2010; Puigserver et al., 1998; Rajakumari et al., 2013; Seale et al., 2009). However, there is a lack of information on the transcription regulators possibly involved in the “plasticity” and/or “trans-differentiation” of white to beige adipocytes. To address this issue, we identified the most expressed set of genes, which encode TFs. Interestingly, in addition to the generic adipogenesis regulators also known to be involved in beige differentiation, such as *Prdm16*, *Pparg*, *Pgc1a*, and *Ppara*, we also showed *Clock* and *Ppara* (both up), for CL and Cold, respectively, and *Egr1* (downregulated) for both treatments. Zinc finger transcription factor EGR1 is a negative regulator of the fat beige program. Loss of Egr1 in mice promotes browning in the absence of external stimulation and leads to increased *Ucp1* expression, which encodes the critical thermogenic mitochondrial uncoupling protein-1 (Milet et al., 2017). Besides, during the trans-differentiation processes, the two treatments seem to activate a different set of genes along their trajectory. This may suggest that, although the endpoint is similar, the possible pathways activated may be different.

***Limitations should be addressed***; our paper show descriptive evidence generated through *in silico* analysis from several angles, but the scope of this study lacks the appropriate *in vivo* models to show a mechanistic link to certain physiological functions. Future studies are needed to validate the metabolic impact of our findings concerning adipose tissue heterogeneity, plasticity, and remodeling. Moreover, secretome and membranome prediction using transcriptomic data as input also needs to be carefully evaluated considering the several mechanisms of transcriptional regulation in mammals. Our workflow template proposes a global analysis of an adipocyte atlas; thus, there is no proper filter from contaminated droplets. The choice of a cutoff may be arbitrary. Finally, the adipocyte trajectory herein described may differ between depots and thermogenic challenges, and further studies are needed to outline and validate the trajectories in different adipocytes subpopulations.

## Conclusion

In summary, we present a multidimensional reanalysis at a single-nucleus resolution, which allows the recovery of nuclei of cell types in adipose tissue. Using this tool, it was possible to show the plasticity of adipocytes, which corroborates with recent data from AT snRNAseq (Rajbhandari et al., 2019; Sun et al., 2020). In addition to the presence of thermogenic adipocytes subpopulation positive for UCP1 (Ad1-*Ucp1*^High^), additional transcriptome analyses allowed us to infer the presence of different secretory functions under thermogenic challenges activated metabolic pathways, especially those related to the futile cycle. On the other hand, Ad1-*Ucp1*^Low^ seems to have a relevant role, both in energy production, through the pathways independent of UCP1, and in its secretory function. This, in turn, proved to be very specific among the heterogeneous adipocyte populations. Regarding the profile of secretory molecules, most are related to proteins that make up EMC, and the highlighted candidates are potentially secreted via classical pathways and exosomes. Understanding this functional plasticity plays an essential role in understanding mature adipocytes’ role as a “central’’ cell type modulating tissue remodeling.

## Supporting information

Figue S4

## Acknowledgments

This work was supported by grants from the NIH DK117161, DK117163 to SRF and P30-DK-046200 to Adipose Biology and Nutrient Metabolism Core of Boston Nutrition and Obesity Research Center, by Sao Paulo Research Foundation (FAPESP) Grants: 2018/20905-1 and 311319/2018-1, the National Council for Scientific and Technological Development, CNPq (282 311319/2018-1 to MLBJr, 311530/2019-2 to RFC, and scholarship #870415/1997-2 to SSC) and by the Coordination for the Improvement of Higher Education Personnel, CAPES (scholarship #88882.378695/2019-01 to CAOBJr)

## Data Statement

The results shown here are based upon data generated by the GSE133486 and the institutional ethical review board approval is not needed.

## Declaration of Interests

The authors declare no conflicts of interest.

## Author Contributions

CAOBJr, performed all R analysis; conceived and wrote all algorithms and the pipeline and participated in the manuscript’s writing; SSC, performed cellular component prediction analysis and participated in the manuscript’s writing; CPA, performed the pseudotime analysis and edited the manuscript; NR, interpreted data and edited the manuscript; WASJr, interpreted edited the manuscript; SRF, design and interpreted edited the manuscript; RFC, was responsible for conceptualization, design, and supervision, MLBJr was responsible for conceptualization, design, and supervision. All authors reviewed and approved the final manuscript.

---

**Fig. S1. Overview of markers and transcriptional profile of adipose tissue cells identified by the new snRNA-Seq template. (A)** *t*-SNE plot of a dataset from Rajbhandari, P *et al*. (2019) distributed by 17 clusters. **(B)** Metacell analysis classification of the “fat cake” 17 clusters in 4 cell types: adipocyte (AD), endothelial (EN), immune (IM), and progenitors (PG). Pie charts show the corresponding percentage. **(C)** SCCAF plot showing the over clustering optimization. **(D)** SCCAF plot showing the Cross-Validation (CV) and test for the different rounds of over clustering. **(E)** Unsupervised markers for each classified cell type generated by the analysis of metacell. **(F)** ORA analysis of DEGs from the four cell types identified. The intensity of the color in the dotplot indicates the enrichment significance by the combined score. Circle sizes correspond to the -log10 adjusted P-value (padj). Each dot plot represents an individual gene set related to the KEGG, WikiPathways, Jensen tissues, and GO biological process

**Fig. S2. Transcriptome-based interactome analysis reveals Ad3-Ad4 cellular interactions.(A)** SCCAF plot showing the over clustering optimization. **(B)** SCCAF plot showing the Cross-Validation (CV) and test for the different rounds of over clustering. **(C)** Violin plot of the canonical mature adipocyte markers. **(D)** Volcano plot representation showing the up and down-regulated DEGs for each mature adipocyte subcluster, with the top 5 up and down-regulated DEGs labeled. **(E)** ORA analysis of DEGs from the four cell types identified. The intensity of the color in the dotplot indicates the enrichment significance by the combined score. Circle sizes correspond to the -log10 adjusted P-value (padj). Each dot plot represents an individual gene set related to the Jensen tissues and Mouse Gene Atlas. **(F)** Pie charts showing the percentage of genes belonging to the secretome, membranome, and other cellular compartments from Ad3 and Ad4 subclusters. **(G)** Heat-scatter plot representing upregulated membranome (upper) and secretome (bottom) components identified in Ad3 and Ad4 subclusters. The color of the circles corresponds to the average logFC. **(H)** Interactome representing cell-cell communication between Ad3 and Ad4 subclusters. P-values are indicated by circle size. The average expression level of interacting molecule 1 in subcluster Ad3/Ad4 and interacting molecule 2 in subcluster Ad4/Ad3 are indicated by color.

**Fig. S3. General analysis of thermogenic treatment in matures adipocyte subpopulation.(A)** Gene-expression heatmap of the DEGs of mature adipocyte nuclei subclusters according to each treatment (Cold, CL, and RT) compared to all others. Genes are represented in rows and cell clusters in columns. **(B)** Selected top categories from ORA analysis of DEGs from mature adipocyte nuclei subclusters according to each treatment (Cold vs. RT and CL vs. RT). The intensity of the color in the dotplot indicates the enrichment significance by the combined score. Circle sizes correspond to the -log10 adjusted P-value (padj). Gene set names are colored according to the GO biological process (purple), Jensen tissues (red), Kyoto Encyclopedia of Genes and Genomes (KEGG, blue), and WikiPathways (orange). **(C)** Heat maps showing the expression of genes related to fatty acid oxidation, tricarboxylic acid cycle, fat acid transport, glycolytic process, triglyceride/fatty acid cycle, and *de novo* lipogenesis in the five mature adipocytes subclusters. **(D)** Venn diagram representing the overlapping between Ad1 and Ad5 subclusters identified in the secretome, membranome, and other cellular locations. **(E)** Pie charts showing the percentage of genes belonging to the secretome, membranome, and other cellular compartments from Ad1 and Ad5 (CL) and Ad2 (Cold). **(F)** Heat-scatter plot representing upregulated membranome (left) and secretome (right) components identified in Ad1 and Ad5 subclusters. The color of the circles corresponds to the average logFC. **(G)** Heat-scatter plot representing upregulated membranome (left) and secretome (right) components identified in Ad2 subcluster. The color of the circles corresponds to the average logFC. **(H)** Interactome representing cell-cell communication between Ad1 and Ad5 subclusters. P-values are indicated by circle size. The average expression level of interacting molecule 1 in subcluster Ad1/Ad5 and interacting molecule 2 in subcluster Ad5/Ad1 is indicated by color. **(I)** Monocle-generated plots presenting pseudotime ordering and differentiation trajectory of CL and RT conditions. The five monocle assigned states are shown. Green background represents the three main thermogenic genes (classic). The yellow background represents the earlier expressed genes in the trajectory. The blue background represents the later expressed genes in the trajectory. The characterized genes are DEGs throughout the trajectory. **(J)** Monocle-generated plots presenting pseudotime ordering and differentiation trajectory of Cold and RT conditions. The seven monocle assigned states are shown. Green background represents the three main thermogenic genes (classic), the yellow background represents the earlier expressed genes in the trajectory, and the blue background represents the later expressed genes in the trajectory. The represented genes are DEGs throughout the trajectory.

**Fig. S4. General characterization of thermogenic main metabolic pathways, secretome prediction, and TF in Ad1-*Ucp1*^High^ and Ad1-*Ucp1*^Low^.(A)** Pie charts show the corresponding percentage of Ad1, Ad2 and Ad5 subclusters expressing the thermogenic genes *Ppara*, *Dio2*, *Prdm16*, *Elovl3*, and *Cox8b*. **(B)** Gene-expression heatmap of all DEGs in Ad1-*Ucp1*^High^ vs. Ad1-*Ucp1*^Low^ comparison. Genes are represented in rows and cell clusters in columns. **(C)** Heat maps showing the expression of genes related to glycolytic process, arginine/creatine and proline metabolism V, and SERCA2 pathways in the Ad1-*Ucp1*^High^ and Ad1-*Ucp1*^Low^. **(D)** Heat-scatter plot representing upregulated membranome (left) and secretome (right) components identified in Ad1-*UCP1*^High^ and Ad1-*UCP1*^Low^. The color of the circles correspond to the average logFC **(E)** Bar plot demonstrating the -log(p-values) of the enriched transcription factors (TFs) in Ad1-*UCP1*^High^ predicted using X2K database. Protein-Protein interactions (PPI) of TFs (green nodes) and targeted genes (grey nodes). The larger the circles, the higher the betweenness centrality value of the node. Gray lines highlight the interactions. Interactions were visualized using Cytoscape v3.7.2. **(F)** Bar plot demonstrating the -log(p-values) of the enriched transcription factors (TFs) in Ad1-*UCP1*^Low^ predicted using the X2K database. Protein-Protein interactions (PPI) of TFs (green nodes) and targeted genes (grey nodes). The larger the circles, the higher the betweenness centrality value of the node. Gray lines highlight the interactions. Interactions were visualized using Cytoscape v3.7.2. **(G)** Heatmap showing the average expression for 42 gene targets from 7 main TFs in Ad1-*UCP1*^High^ cells (green background) and 13 gene targets from 5 main TFs in Ad1-*UCP1*^Low^ cells (yellow background). These targets and TFs are related to the DEGs from each comparison through the Transcription Factor Enrichment Analysis (TFEA) using the X2K database.

## STAR Methods Key resources table

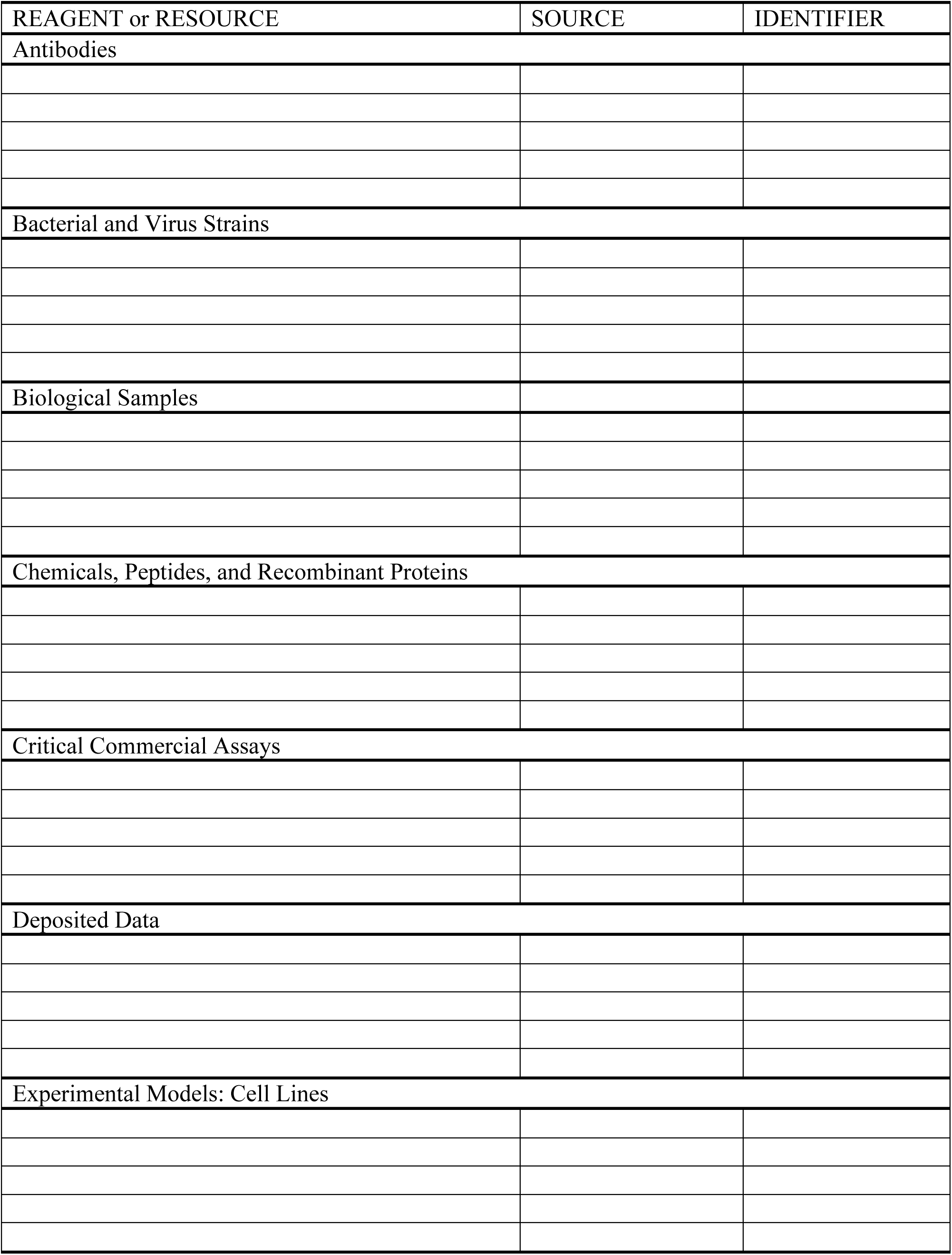
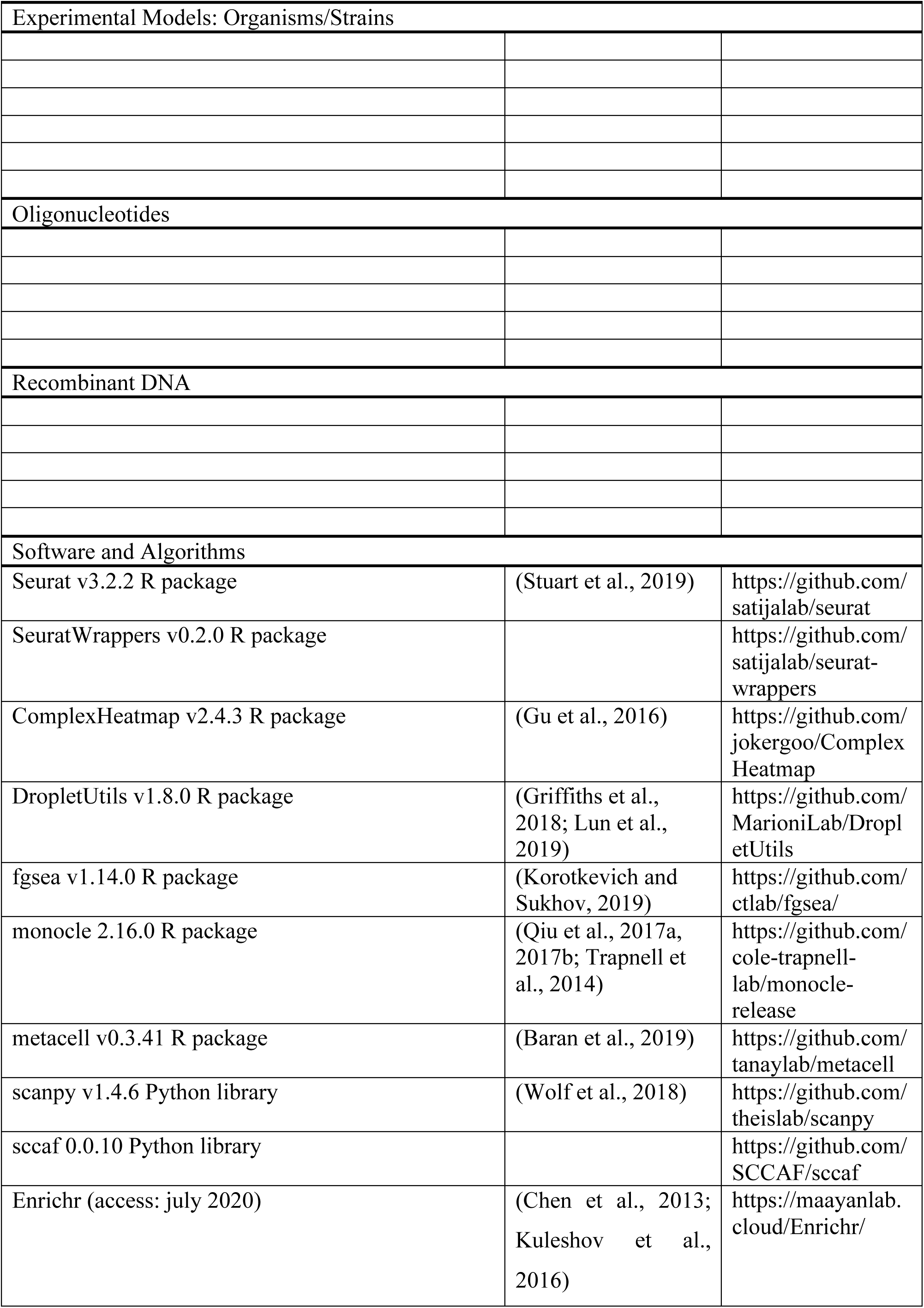
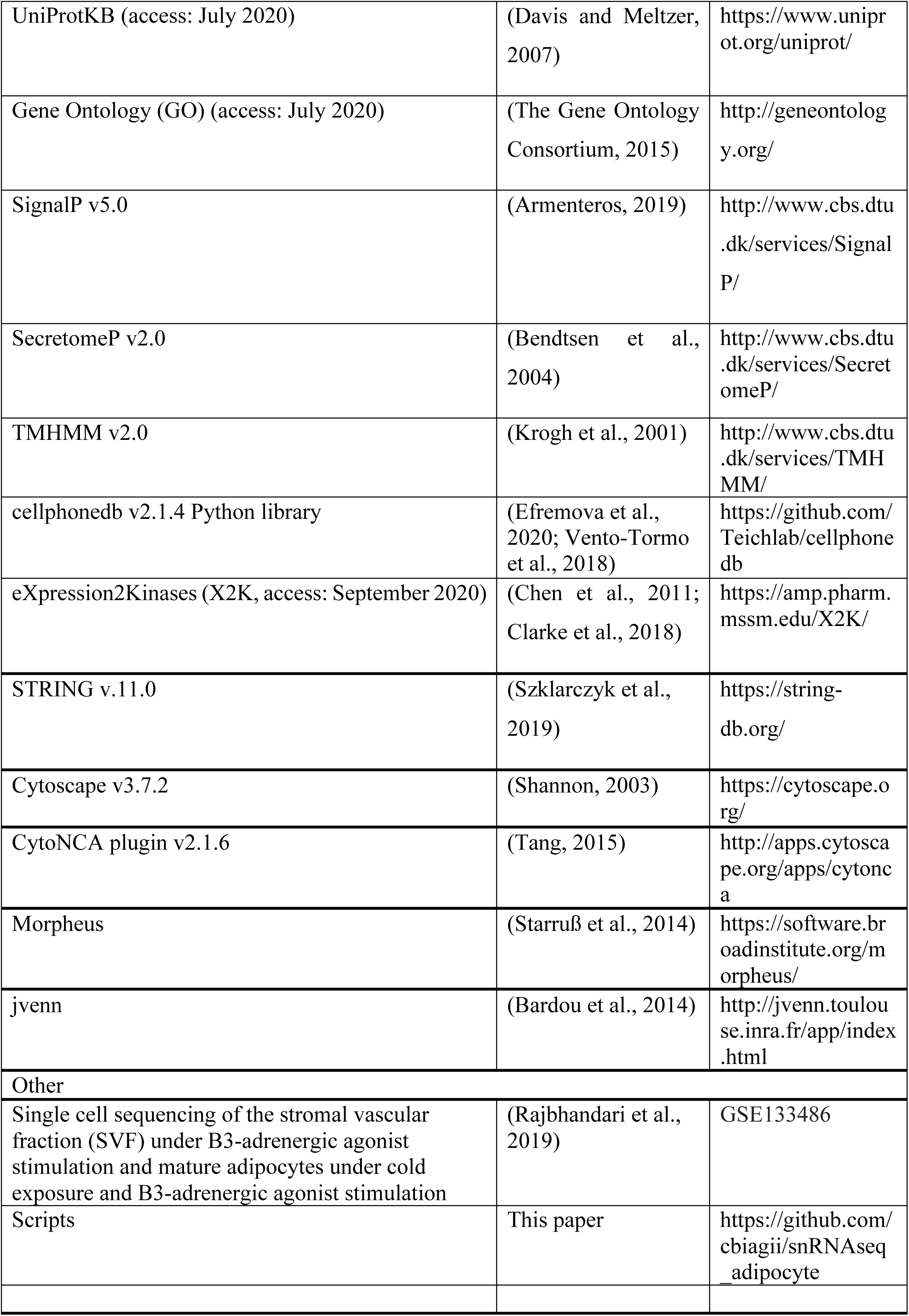

## Methods Details

### Single-cell RNA-seq data

The scRNA-seq data from mature adipocytes under cold-challenge and B3-adrenergic agonist stimulation were acquired from the Gene Expression Omnibus (GEO) database under the series number GSE133486 (Rajbhandari et al., 2019), which contains 6 data of mouse SVF and 10 data of mouse adipose nuclei generated using Drop-Seq and 3’ V3 chemistry kit on Chromium Single-cell controller (10x Genomics), respectively.

### Data pre-processing

The filtered feature-barcode matrix was used in the following analysis. All additional analysis were performed using Seurat v3 (Butler et al., 2018; Stuart et al., 2019). First, to reproduce the results obtained by Rajbhandari et al., the same procedures described in the paper were used. For the single nuclei data reanalysis, only data of mouse adipose nuclei wild-type mice underwent cold-challenge and B3-adrenergic agonist stimulation were used. Nuclei with less than 200 and more significant than 3000 genes detected, more than 10% of mitochondrial percentage genes, and P condition were excluded from the analysis. We assigned scores for S and G2/M cell cycle phases based on previously defined gene sets (Tirosh et al., 2016) using the CellCycleScoring function for clustering of all cells. The regularized negative binomial regression was used to normalize UMI count data using the sctransform workflow (Hafemeister and Satija, 2019), regressing out against the number of UMIs per cell, S phase score, and G2/M phase score. Scaled data was used as an input into PCA based on variable genes. Clusters were identified using Shared Nearest Neighbor (SNN) based clustering based on the first 26 PCs (corresponds to a PCA cumulative proportion greater than 80%) and resolution = 1. The same principal components were used to generate the t-SNE projection, which was developed with a maximum of 2000 iterations.

### Optimal number of clusters

To find the optimal number of clusters, the SCCAF was used with an accuracy threshold of 80%. The clustering calculated previously was used as initial clusters, and the h5ad file used as input was generated using the SeuratToH5ad function. The optimize, skip-assessment, produce-rounds-summary, and optimisation-plots-output parameters were used in this first step to use the sccaf command to round optimization. Then the sccaf-assess command was used to determine the round to be used as a final result by through the observation of accuracies for each round on multiple iterations. For this step the default parameters were used using 20 iterations. Finally, the last step is to generate a plot used to compare the accuracy between different rounds. For this purpose, the sccaf-assess-merger command was employed using the results from step one and two.

### Cell-type classification

To classify the cell types Metacell (Baran et al., 2019) was used with the default parameters. Firstly, we used some initial markers as *Adrb3* for Adipocyte, *Pecam* for Endothelial, *Ptprc,* and *Cd19* for Immune and *Cd34,* and *Pdgfra* for Progenitor cell type. Based on these markers and cell types, the tool returned a list of new markers that can separate these cell types. To select these new markers, we performed isolation of each cell type group to check each marker’s expression. The markers that had an expression in the given group, greater than 90% of the quantile, were considered markers for the respective group. These are the markers: *Acsl1*, *Plin4*, *Mlxipl*, *Pck1,* and *Adrb3* for Adipocyte, *Btnl9*, *Ushbp1*, *Egfl7*, *Mcf2l* and *Ptprb* for Endothelial, *Zeb2*, *Trps1*, *Runx1*, *Ptprc,* and *Adap2* for Immune, *Dcn*, *Celf2*, *Meg3*, *Col1a2* and *Col3a1* for Progenitor cell types. So, based on these markers, the tool was able to classify the cell types.

### Optimal mature adipocyte subclusters

The mature adipocyte cluster was subsetted using an optimal number of clusters the SCCAF (described above) was used with an accuracy threshold of 80%. RT (Ad3 e Ad4), Cold (Ad1 e Ad2), CL (Ad1 e Ad5); adipocyte clusters of cells (Ad1, Ad2, Ad3, Ad4 and Ad5)

### Differential Expression and Enrichment analysis

Differentially expressed genes between the different conditions each cluster/cell type were identified using FDR < 0.05 and/or |avg_logFC| > 0.25. Functional enrichment analysis was performed using the Enrichr tool (Chen et al., 2013; Kuleshov et al., 2016). For better visualization of the data, the Adaptively-thresholded Low Rank Approximation (ALRA) (Linderman et al., 2018) imputation method and Nebulosa (Kernel Gene-Weighted Density Estimation) (Jose Alquicira-Hernandez, 2020) was used.

### Transcriptome-based secretome analysis

The differentially expressed genes in the mature adipocyte subclusters (Ad1, Ad2, Ad3, Ad4, and Ad5) were filtered for genes encoding secreted proteins based on a pipeline of four databases and tools. UniProtKB (The UniProt Consortium, 2019) annotation of subcellular localization was accessed to select proteins classified as “Secreted” and Gene Ontology (GO) (The Gene Ontology Consortium, 2015) annotation of the cellular component was used for selection of “Extracellular” proteins. To confirm those results, the combined lists of proteins generated by UniprotKB and GO were analyzed using the algorithms SignalP 5.0 (Roh et al., 2018), SecretomeP 2.0 (Bendtsen et al., 2004). SignalP server (Armenteros, 2019) was used to identify classical secretory proteins (presenting signal peptide considering the D-value >0.45). Proteins without signal peptide were evaluated in the SecretomeP 2.0 server to determine non-classical secreted proteins, using the cutoff for a neural network (NN) score >0.6. The same strategy was used for clusters of adipocytes highly expressing *Ucp1* compared to adipocytes with lower levels of *Ucp1*. Each condition’s predicted secretome was visualized using heat-dot plot using Morpheus software (Starruß et al., 2014). We also verified the secretion via exosomes by accessing the Exocarta database (http://www.exocarta.org/) (Simpson et al., 2012).

### Membranome prediction

The differentially expressed genes in the mature adipocyte subclusters (Ad1, Ad3, Ad4, and Ad5) were filtered for genes encoding cell membrane proteins. UniprotKB and GO were also used for the membranome annotation filtering in the proteins classified at “cell membrane” and “plasma membrane”, respectively. The list generated was confirmed using TMHMM 2.0 (Krogh et al., 2001) algorithm selecting only proteins with predicted number of transmembrane helices (PredHel) greater than 1. This prediction analysis classifies transmembrane proteins without discriminating if the protein is located on plasma or vesicles or organelles membranes. Thus, we manually reviewed the literature of our selected top 5 genes of interest to confirm which membrane they belong to. The same strategy was used for subclusters of adipocytes highly expressing *Ucp1* compared to adipocytes with lower levels of *Ucp1*.

### Cell-cell interaction

We used the computational framework CellPhoneDB to predict cell-cell communication using its repository of curated ligand-receptor interactions for single-cell transcriptomic data (Efremova et al., 2020; Vento-Tormo et al., 2018). We used the default setting to select the statistically relevant interaction (p-value < 0.05) between the mature adipocyte subclusters belonging to RT group (Ad3 vs. Ad4) and the CL group (Ad1 vs. Ad5).

### Pseudotime analysis

To analyze the trajectory development of adipocyte clusters, an unsupervised pseudo temporal analysis was performed using Monocle2 (Qiu et al., 2017a, 2017b; Trapnell et al., 2014). The Seurat object with cluster information was extracted and converted to a Monocle2 CellDataSet. Monocle2 uses DDRTree, a reversed graph embedding algorithm to predict biological trajectories, to reduce the high-dimensional scRNA-seq data space and predict how cells progress through a given biological process based on global gene expression levels. Monocle2 offers ideal unsupervised pseudotime analysis for this study as it indicates branch points and trajectory states without cell fate input information. Following size factor, dispersion estimates, trajectory ordering genes were called by testing the differential expression of genes expressed with min_expr = 0.1 in >= 10 cells against the 5 clusters of adipocytes, selecting the genes that have qval < 0.01. Data dimensionality was reduced using the reduceDimension function with max_components set to 2 and reduction_method set to DDRTree. The cells were ordered according to the state that represents the initial condition (state 1). DEGs across pseudotime were determined using the differentialGeneTest function filtering by qval < 0.01. Resultant genes were ordered by q value, and the top-500 genes changing in pseudotime were visualized using the plot_pseudotime_heatmap function.

### Transcription factors (TFs) enrichment analysis and Protein-protein interaction (PPI)

We used the eXpression2Kinases (X2K) (Chen et al., 2011; Clarke et al., 2018) workflow to identify the upstream TFs of the DEGs within subclusters with high and low gene expression of *Ucp1*. We selected the enriched TFs (p-value < 0.05) to construct the PPI network with their targeted genes found as DEGs in the adipose single-cells expressing or not *Ucp1* in cluster Ad1. PPI networks were conducted using STRING v.11.0 (https://string-db.org/). Only medium confidence interactions were included (interaction score of at least 0.4), and the disconnected nodes were omitted in the network. Visualization and data annotation of PPI networks were constructed using Cytoscape v3.7.2(Shannon, 2003). The CytoNCA plugin (Tang, 2015) was used to calculate the betweenness centrality values of each node.

### Data representation

Heat-scatter plot was generated using Morpheus (Starruß et al., 2014) (https://software.broadinstitute.org/morpheus). Venn diagrams were plotted using the web server jvenn (Bardou et al., 2014).

### Data and code availability

The accession number for the single-nuclei sequencing data from Rajbhandari *et al*. (2019) reported in this paper is GEO: GSE133486. All analysis code is available on GitHub at https://github.com/cbiagii/snRNAseq_adipocyte.

## Notes

### Competing Interest Statement

The authors have declared no competing interest.

### Summary of Updates

Introduction session and references were updated; author affiliations updated.

