## Supplementary figures and images for "Multidimensional Single-Nuclei RNA-Seq Reconstruction of Adipose Tissue Reveals Adipocyte Plasticity Underlying Thermogenic Response"

### Figue S4

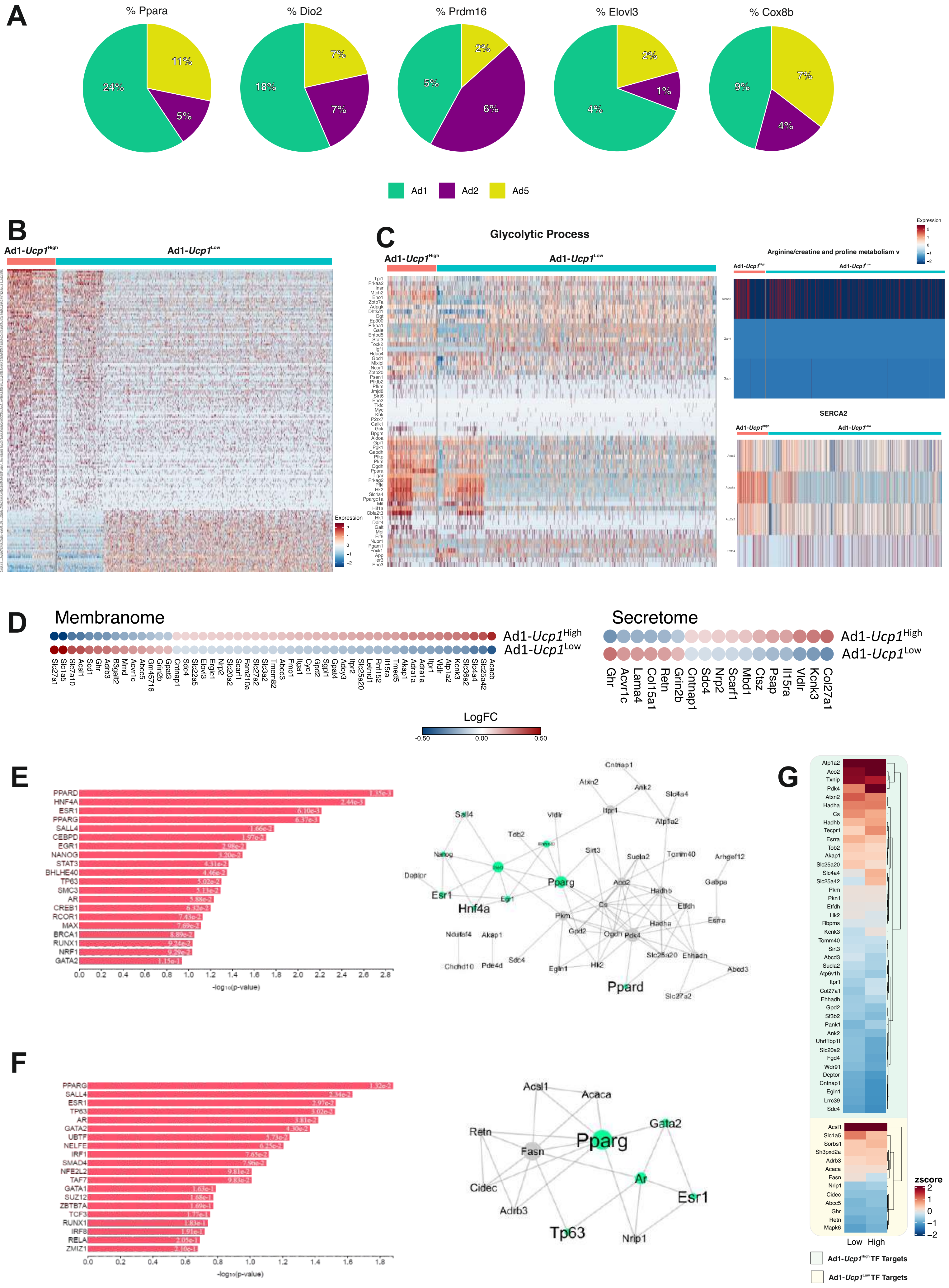

### Figure S1

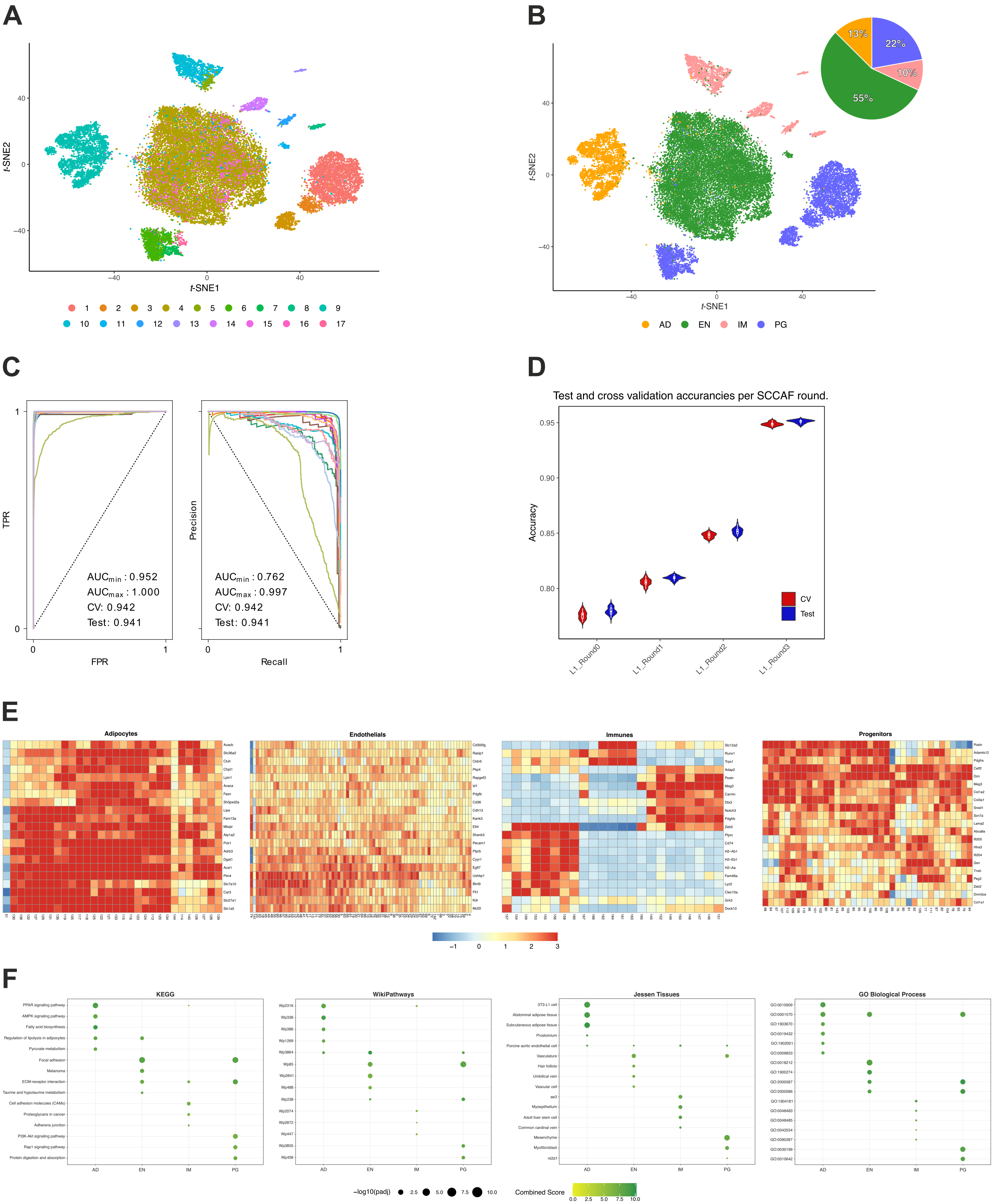

### Figure S2

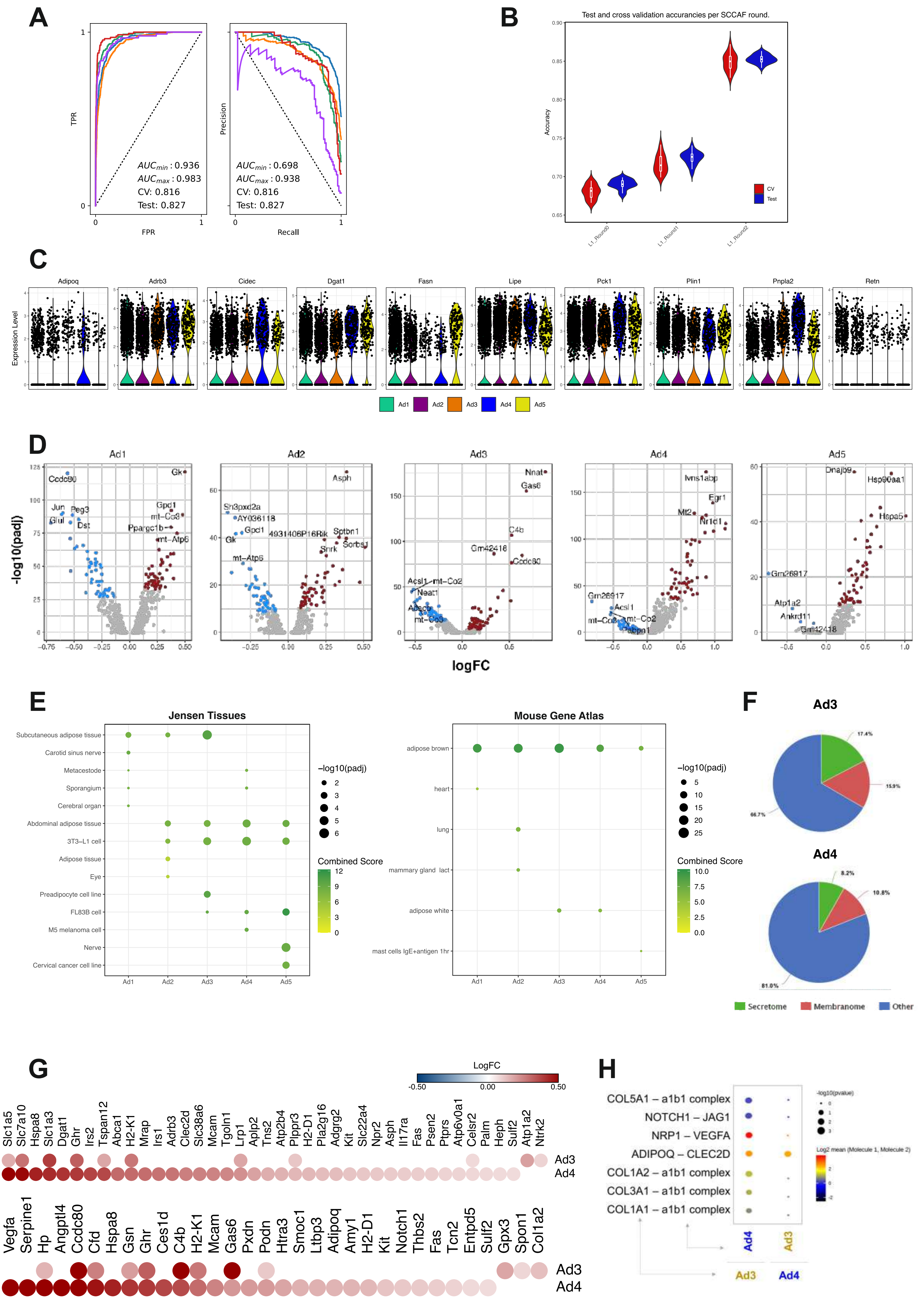

### Figure S3

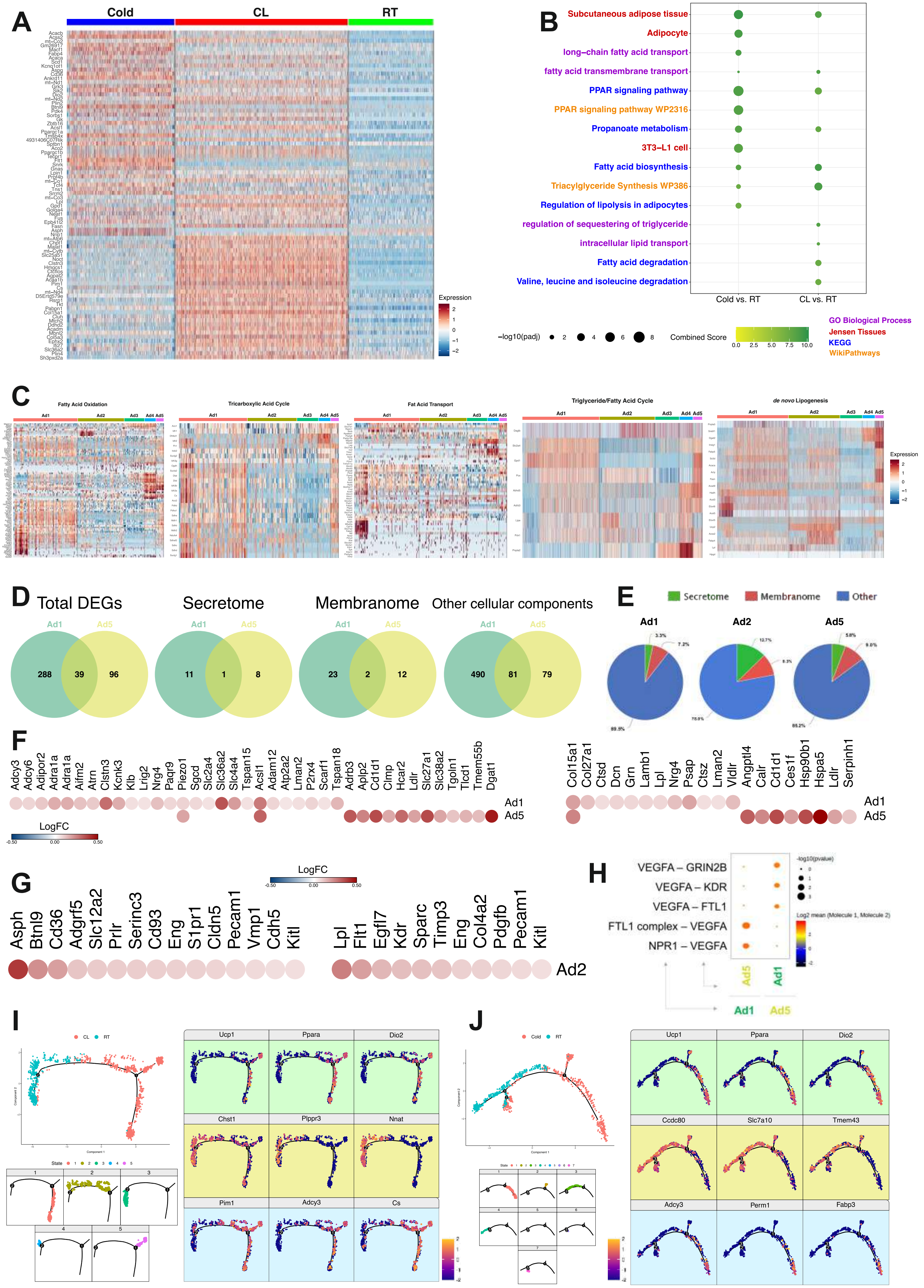
